# Cytokinin response induces immunity and fungal pathogen resistance in tomato by modulating cellular trafficking of PRRs

**DOI:** 10.1101/2020.03.23.003822

**Authors:** Rupali Gupta, Lorena Pizarro, Meirav Leibman-Markus, Iftah Marash, Maya Bar

**Author notes:** Corresponding author: Dr. Maya Bar.

## Abstract

Plant immunity is often defined by the “immunity hormones”: salicylic acid (SA), jasmonic acid (JA), and ethylene (ET). These hormones are well known for differentially regulating defense responses against pathogens. In recent years, the involvement of other plant growth hormones such as auxin, gibberellic acid, abscisic acid, and cytokinins (CKs) in biotic stresses has been recognized. Previous reports have indicated that endogenous and exogenous CK treatment can result in pathogen resistance. We show here that CK induces systemic tomato immunity, modulating cellular trafficking of the PRR LeEIX2 and promoting biotrophic and necrotrophic pathogen resistance in an SA and ET dependent mechanism. CK perception within the host underlies its protective effect. Our results support the notion that CK acts as a priming agent, promoting pathogen resistance by inducing immunity in the host.

## Introduction

During plant-pathogen interactions, plants must sense and respond to many signals to balance growth and defense in an adverse environment (Jones and Dangl, 2006). The response to external stimuli through cellular signaling events can lead to resistance or susceptibility to pathogens, depending on the environment, pathogen virulence factors, and the genetics of the host plant.

Plant immune responses have evolved sophisticated strategies to suppress pathogen infection, and the two major layers of which are pathogen-associated molecular patterns (PAMPs)-triggered immunity (PTI) and effector-triggered immunity (ETI) (Spoel and Dong, 2012). These phenomena reflect the dynamic balance between the ability of the plant to activate defense responses against pathogen and the capability of the pathogen to suppress the plant’s immune system (Dodds and Rathjen, 2010). Plant induced immunity has become an interesting area to study the perception of pathogens and activation of host defense regulatory mechanisms (Macho and Zipfel, 2014). The induced immunity of plants displaying increased resistance is often not attributed to direct activation of defenses, but, rather, to a rapid, stronger activation of basal defense signaling upon exposure to pathogens.

Plant immunity is often defined by what are considered the “immunity hormones”: salicylic acid (SA), jasmonic acid (JA), and ethylene (ET). These hormones are well known for differentially regulating defense responses against pathogens (Bari and Jones, 2009; Hatsugai et al., 2017). In recent years, the involvement of other, more “classical” plant growth hormones such as auxin, gibberellic acid (GA), abscisic acid (ABA), and cytokinins (CKs) in biotic stresses has been recognized (Shigenaga et al., 2017; Chanclud and Morel, 2016).

Cytokinin (CK) is an important developmental regulator, having activities in many aspects of plant life and its response to the environment. CKs are involved in diverse processes including stem-cell control, vascular differentiation, chloroplast biogenesis, seed development, growth and branching of root, shoot and inflorescence, leaf senescence, nutrient balance and stress tolerance (Muller and Sheen, 2007). The roles of CK in plant growth and development have been reviewed extensively (Werner and Schmülling, 2009; Sakakibara, 2006; Keshishian and Rashotte, 2015; Mok and Mok, 2001).

In some cases, plant pathogens can secrete CKs, or induce CK production in the host plant. The hemibiotrophic actinomycete *Rhodococcus fascians* produces CKs. Recognition of *R. fascians* derived CKs is essential for symptom development in Arabidopsis (Pertry et al., 2009). The spores of biotrophic rust and powdery mildew fungi contain CKs, which may be associated with green islands at the infection sites (Király, Z., El Hammady, M., Pozsár and Kiraly, Z., Hammady, M.E., and Pozsar, 1967; Kiraly, Z., Pozsar, B., and Hammady, 1966). It has been suggested that to achieve pathogenesis in the host, CK-secreting biotrophs or hemibiotrophs manipulate CK signaling to regulate the host cell cycle and nutrient allocation (Jameson, 2000).

Works concerning the roles of CK in the plants’ interaction with microbes that do not produce CK are less abundant. High levels of CKs were found to increase the plants’ resistance to some viral pathogens and herbivores (Ballaré, 2011). Transgenic overexpression of CK-producing IPT genes increased Arabidopsis resistance to *Pseudomonas* (Choi et al., 2010), while overexpression of genes encoding CK oxidase, or mutating the endogenous CK AHK receptors, enhanced Arabidopsis pathogen susceptibility (Choi et al., 2011, 2010; Argueso et al., 2012). In another study, CKs were found to mediate enhanced resistance to *Pseudomonas syringae* in tobacco (Grosskinsky et al., 2011). Different mechanisms have been suggested for this enhanced resistance. In Arabidopsis, it was suggested that CK-mediated resistance functions through SA dependent mechanisms, based on the finding that ARR2,(a positive regulator of CK signaling) interacts with TGA3 (a transcription factor involved in inducing SA-responsive genes) in the regulation of the disease marker gene PR1 against biotrophic infections in plants (Choi et al., 2010). An additional study suggested that CK signaling enhances the contribution of SA-mediated immunity in hormone disease networks (Naseem et al., 2012). CK was proposed to function in Arabidopsis immunity against the biotroph *H. arabidopsidis* through repression of type-A response regulators (Argueso et al., 2012). In tobacco, an SA-Independent, phytoalexin-dependent mechanism was suggested (Grosskinsky et al., 2011).

In this work, we investigated the effects of CK on disease resistance and immunity in tomato, demonstrating that CK ameliorates disease outcomes of the tomato necrotrophic fungus *Botrytis cinerea* and biotrophic fungus *Oidium neolycopersici*, that CK activates tomato immunity, and that CK signaling is activated in tomato in response to *B. cinerea*. We show that high CK levels activate the plant defense machinery, and that CK response within the plant serves as a systemic immunity signal. CK promotes PRR trafficking and requires SA and ET, but not JA mechanisms, to exert its full effect in tomato defense.

## Results

### Exogenous CK treatment ameliorates tomato disease

Works describing the role of CK in plant disease were obtained only in select plant-pathogen experimental systems (Albrecht and Argueso, 2017). To investigate the role of CK in tomato fungal disease response, we examined the effect of CK on pathogenesis of a necrotrophic fungal pathogen, *B. cinerea* (*Bc*), which is the causative agent of grey mold in over 1000 different plant hosts, and a biotrophic fungal pathogen, *O. neolycopersici* (On), a causative agent of powdery mildew disease. The results are presented in Figure 1.

**Figure 1.**
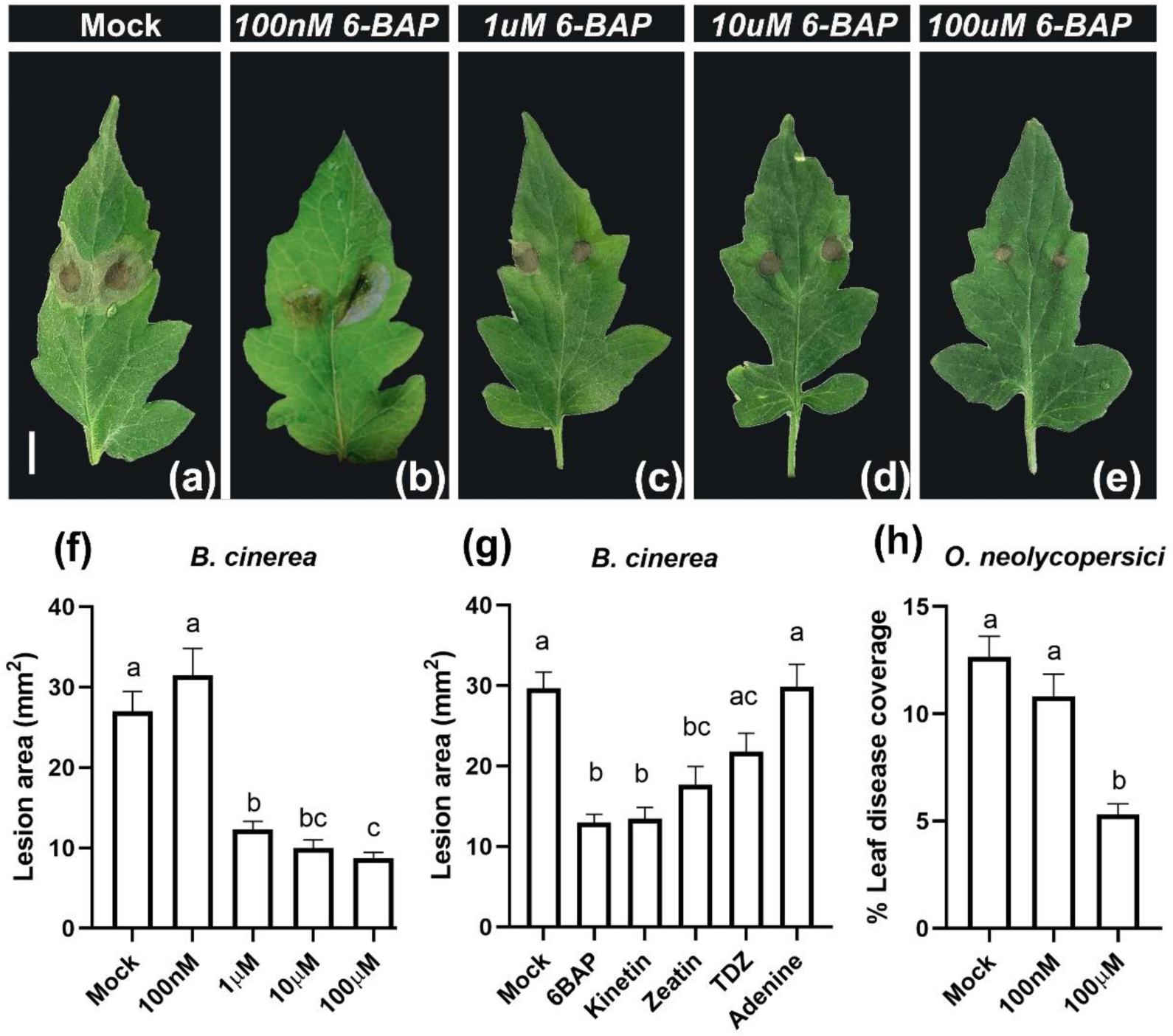
CK reduces disease symptoms of necrotophic and biotrophic fungi in tomato. (a)-(f) *S.lycopersicum cv* M82 plants were spray-treated with indicated concentrations of 6-Benzylaminopurine (6-BAP) dissolved in 1uM NaOH, and inoculated with 10 ul of *B. cinerea* spore solution (10^6^ spores ml^-1^) 24 hours later. Bar, 1cm. (f) Lesion area was measured on whole plants 5-7 days after *B. cinerea* inoculation using ImageJ. (g) *S.lycopersicum cv* M82 leaves were treated with 100uM of indicated CK compounds or the control adenine, and, after 24 hours, detached from the plant and inoculated with 10 ul of *B. cinerea* spore solution (10^6^ spores ml^-1^). Lesion area was measured 5-7 days after *B. cinerea* inoculation using ImageJ. Similar disease levels were achieved with *B.cinerea* on whole plants (f) and detached leaves (g). (h) *S.lycopersicum cv* M82 plants were treated with indicated concentrations of 6-BAP and spray inoculated with *O. neolycopersici* (10^4^ ml^-1^ spores) 24 hours later. Leaf disease coverage was calculated ten days after *O. neolycopersici* inoculation. In all cases, plants treated with 1 uM NaOH were used as mock. Graphs represent the results of at least 4 independent experiments ±SEM. Results were analyzed for statistical significance using one-way ANOVA with a Dunnett post hoc test. Different letters indicate statistically significant differences. (f) N≥40, p<0.0001. (g) N≥30, p<0.0001. (h) N≥9, p<0.0001.

Wild type (WT) *Solanum lycopersicum cv* M82 tomato plants were treated with varying concentrations of the CK 6-Benzyl Amino Purine (6BAP, BA) prior to pathogen infection, and the dose response of disease progression was measured as described in the methodology section. Disease was assessed 5-10 days after pathogen inoculation. CK pre-treatment significantly decreased disease levels of the necrotrophic fungal pathogens *B. cinerea* (*B. cinerea, Bc*), (Figure 1 a-f), and the biotrophic fungal pathogen *Oidium neolycopersici* (Figure 1h). The strength of the disease follows a dose-response to the CK concentration applied.

To test whether CK pre-treatment of additional CK compounds has a similar effect, we examined *B. cinerea* disease progression after pre-treatment with kinetin, *trans*-zeatin, and thidiazuron (TDZ), as well as adenine as a structurally similar control compound. Figure 1g demonstrates that all assayed CKs ameliorate *Bc* disease outcomes, with the exception of TDZ, a phenylurea-derived artificial cytokinin, which is structurally unrelated to the purine-type cytokinins, though it is known to bind strongly to the Arabidopsis CK receptors AHK3 and AHK4 (Romanov et al., 2006). Adenine, the control compound, has no significant effect on disease progression.

### Increased endogenous CK quantity or sensitivity improves tomato disease outcomes

To examine whether endogenous CK levels or response might have a similar effect, WT and CK tomato mutants were assessed for disease resistance or sensitivity in a similar manner. Figure 2 demonstrates that plants with elevated endogenous levels of CK (*pBLS>>IPT7* (Shani et al., 2010) or increased CK sensitivity (*e2522 clausa* (Bar et al., 2016) have significantly lower disease symptoms with *B. cinerea* (Figure 2a-e) and n *O. neolycopersici* (Figure 2g). *pFIL>>CKX4* (Shani et al., 2010), which constantly breaks down its endogenous CK, had significantly increased disease levels in both cases (Figure 2e,g).

**Figure 2.**
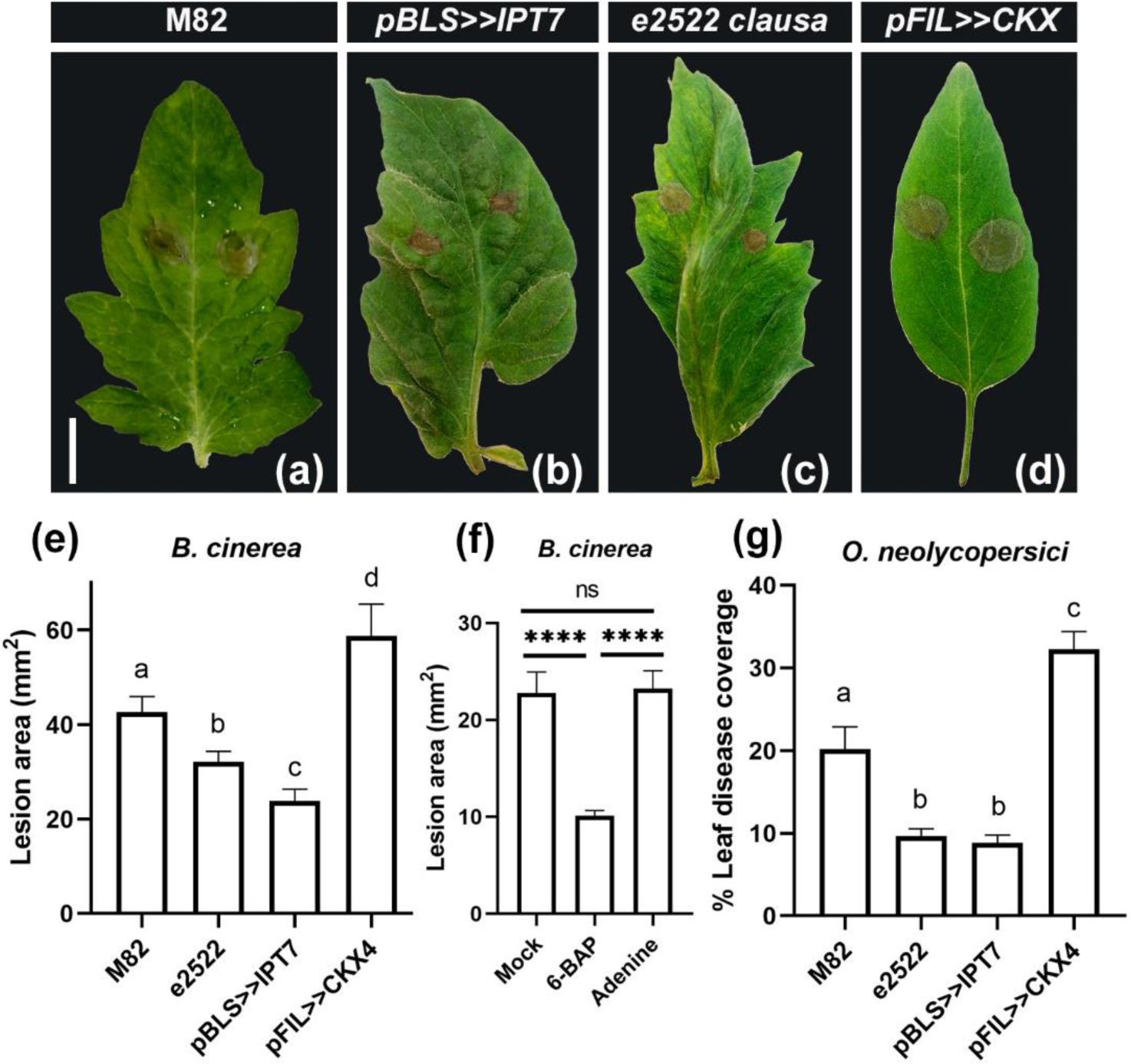
**CK induced disease resistance in tomato depends on host signaling** (a)-(e): *pBLS>>IPT7* (increased cytokinin content transgenic line), *clausa* (increased cytokinin-sensitivity mutant), and *pFIL>>CKX4* (reduced cytokinin content transgenic line) plants, all in a *S.lycopersicum cv* M82 background, were sprayed with 100 uM 6-Benzylaminopurine (6-BAP) dissolved in 1uM NaOH 24 hours before application of a10 ul of *B. cinerea* (*Bc*) spore solution (10^6^ spores ml^-1^). Bar, 1 cm (e) Lesion area was measured 5-7 days after inoculation using ImageJ. (f) *S.lycopersicum cv* M82 plants were soil drenched with 100 uM 6-BAP dissolved in 1uM NaOH or with 100 uM Adenine, and inoculated with 10 ul *Bc* spore solution (10^6^ spores ml^-1^). Lesion area was measured 5-7 days after *Bc* inoculation using ImageJ. (g) *S.lycopersicum cv* M82 plants were sprayed with *O. neolycopersici* (10^4^ ml^-1^ spores) 24 hours after 100 uM 6-BAP treatment. Leaf disease coverage was calculated ten days after *O. neolycopersici* inoculation using ImageJ. In all cases, plants treated with 1 uM NaOH were used as mock. Graphs represent the results of 3-6 independent experiments ±SEM. Results were analyzed for statistical significance using one-way ANOVA with a Dunnett post hoc test. Different letters indicate statistically significant differences. (e) N≥32, p<0.0001. (f) N≥40, p<0.0001. (g) N≥10, p<0.0001.

Modulating CK levels both exogenously (Figure 1) and endogenously (Figure 2) improved tomato disease outcomes. To examine whether this is a systemic effect, CK was also applied by soil drench to the roots, with similar results in *Bc* disease resistance in leaves (Figure 2f), indicating that CK affects tomato disease resistance systemically.

### CK disease amelioration is ET and SA dependent- and JA independent

It was previously reported that CK influences biotrophic disease resistance through regulation of SA in Arabidopsis (Choi et al., 2010; Naseem et al., 2012; Naseem and Dandekar, 2012). We found here that CK induces resistance to necrotrophic pathogens in tomato. Since JA and ET are known to be involved in the response to necrotrophic pathogens (Thomma et al., 1998), we explored the involvement of SA, JA and ET in the amelioration of disease outcomes by CK, conducting pathogenesis assays in SA, JA and ET signaling/biosynthesis tomato mutants. Figure 3 shows that the SA deficient *NahG* transgenic line (Brading et al., 2000) (Figure 3b,h,m), and the ET reduced sensitivity mutant *Never ripe* (*Nr)* (Lashbrook et al., 1998) (Figure 3d,j,m) have no significant *B. cinerea* disease amelioration upon CK treatment. However, the JA insensitive *jai-1* mutant (Li et al., 2002) responds to CK with disease reduction (Figure 3f,l,m). *B. cinerea* disease levels in these mutants without CK treatment matched those known in the literature when compared to their background genotypes: a moderate decrease in *NahG* (Mehari et al., 2015), similar levels in *Nr* (Mehari et al., 2015), and increased levels in *jai-1* (AbuQamar et al., 2008), when compared to the background cultivars (Figure 3m).

**Figure 3.**
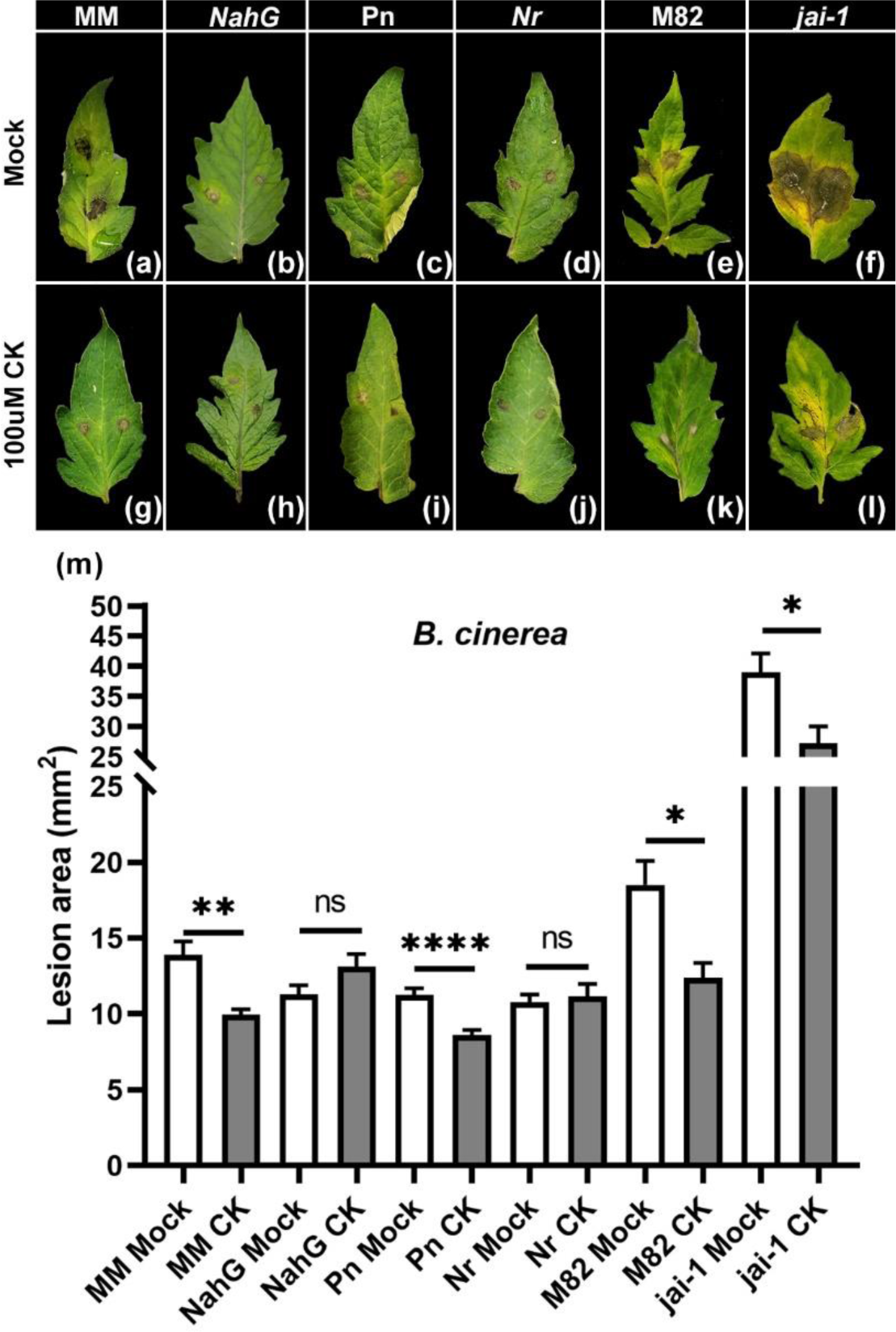
Cytokinin induced disease resistance requires salicylic acid (SA) and ethylene (ET), but not jasmonic acid (JA) signaling. (a)-(m): Leaves of the SA deficient *NahG* and its background line Moneymaker (MM), the Ethylene insensitive Never-Ripe (*Nr*) and its background line Pearson (Pn), and the Jasmonate insensitive *(jai-1*) and its background M82, were spray treated with 100uM cytokinin (CK; 6-benzylaminopurine, 6BAP) dissolved in 1uM NaOH, and, after 24 hours, detached from the plant and inoculated with 10 ul of *B. cinerea* (*Bc*) spore solution (10^6^ spores ml^-1^). (m) Lesion area was measured 5-7 days after inoculation using ImageJ. Plants treated with 1 uM NaOH were used as mock. (m) Graph represents 3-5 independent experiments, N≥17 ±SEM. Results were analyzed for statistical significance using one-way ANOVA with a Dunnett post hoc test, p<0.0001; *p<0.05; **p<0.01; ****p<0.0001..

Our results match the literature for Arabidopsis, demonstrating that CK does not ameliorate *Pst* disease outcomes in a SA deficient background (Choi et al., 2010).

### Altering CK response changes SA profiles upon pathogen infection

Our results indicated that CK induced pathogen resistance in tomato requires the SA pathway. To examine this further, we quantified SA in mock and CK pretreated tissues, as well as in genotypes with altered CK levels/ response, in *B. cinerea* infected and uninfected tomato plants. CK pre-treatment caused changes in SA content. However, the baseline levels of SA in genotypes with altered levels of CK or CK sensitivity-resembled those of the background line (Figure 4). *B. cinerea* inoculation results in a reduction in SA content after 48 hours, External CK pre-treatment or increased endogenous CK sensitivity in the *clausa* mutant both maintain the SA reduction following *Bc* inoculation, though to a significantly lesser degree. Endogenously manipulating CK levels in *pBLS>>IPT7* or *pFIL>>CKX* abolishes the reduction of endogenous SA following *Bc* application. (Figure 4).

**Figure 4.**
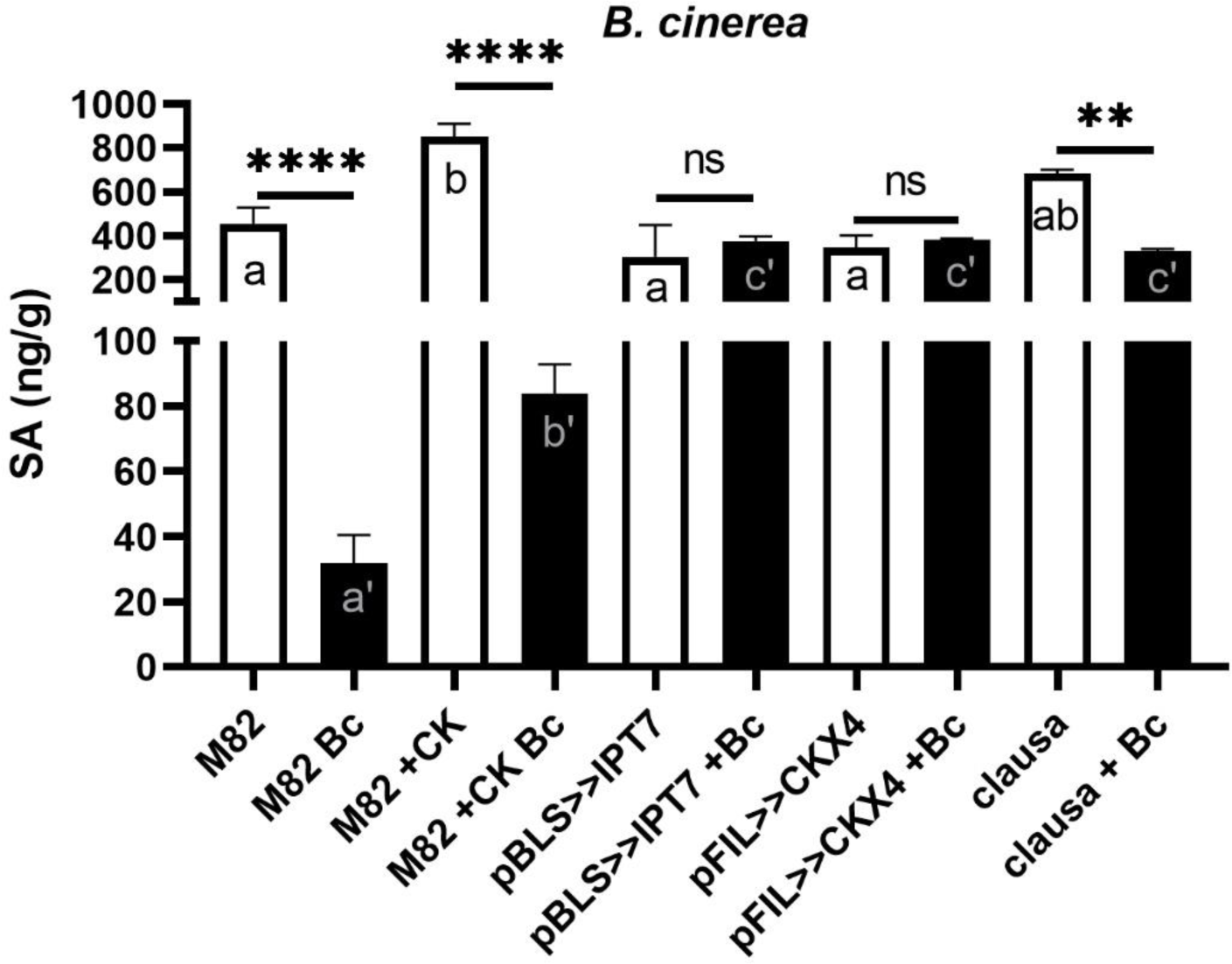
Salicylic acid (SA) content does not directly correlate with cytokinin induced disease resistance. Levels of SA in the indicated genotypes, all in the M82 background, in mock (1 uM NaOH) or CK (100uM 6-Benzylaminopurine, dissolved in 1 uM NaOH) treated plants, and 48 hours after *B. cinerea* (*Bc*) inoculation, were quantified using LC–MS-MS. Average ±SEM presented in all cases, N=3-9. Asterisks and different letters indicate significance in a one-way ANOVA with a Tukey post hoc test, p<0.0001. Asterisks indicate significance in the reduction in SA content 48 hours after *Bc* inoculation. Lower case letters in the white bars indicate significant differences in the SA levels in mock plants. Lower case tagged letters in the black bars indicate significant differences in the SA levels in the different genotypes following *Bc* inoculation. All assays were conducted on intact plants, with leaves harvested for processing and analysis 48 hours after *Bc* inoculation.

### CK induces tomato immunity

Our results indicate that tomato pathogen resistance is modulated by both endogenous and exogenous CK, as was previously reported for *Pst* in Arabidopsis (Choi et al., 2010). Exogenous CK application primes Arabidopsis defense (Albrecht and Argueso, 2017). To examine whether the decrease in fungal disease in the presence of elevated CK levels is paired with increased plant defense in tomato, we tested known hallmarks of immune system activation: ethylene production, ion leakage and ROS. Treating WT M82 plants with exogenous CK results in an increase in ethylene production and conductivity (Figure 5 a-b). In addition to 6BAP, we tested kinetin, *trans*-zeatin, and thiodiazurone (TDZ). Adenine served as a negative control. Kinetin and trans-Zeatin had similar activity as 6-BAP in the activation of plant defenses. TDZ had no effect on ethylene production, and a lower effect on conductivity than the other CKs (Figure 5c-d). Adenine has no significant effect on plant immune responses. We examined tomato genotypes with altered CK levels or response. Genotypes with elevated CK sensitivity or levels have elevated defense responses: the *clausa* mutant and the overexpressor of IPT both have elevated basal levels of ethylene when compared with M82 plants, and IPT also has elevated levels of conductivity (Figure 5 e-f). The defense response mutants *Nr* and *jai-1* responded to CK with ethylene production and ion leakage at similar levels to those of their respective background cultivars, Pn and M82 (Figure 6a,b). The SA deficient *NahG* had reduced ethylene production in response to CK when compared with its background cultivar MM, and, unlike MM, did not respond to CK with an increase in ion leakage (Figure 6a,b). Baseline defense responses without CK were not significantly different between the mutant genotypes and their background lines (Figure 6a,b).

**Figure 5.**
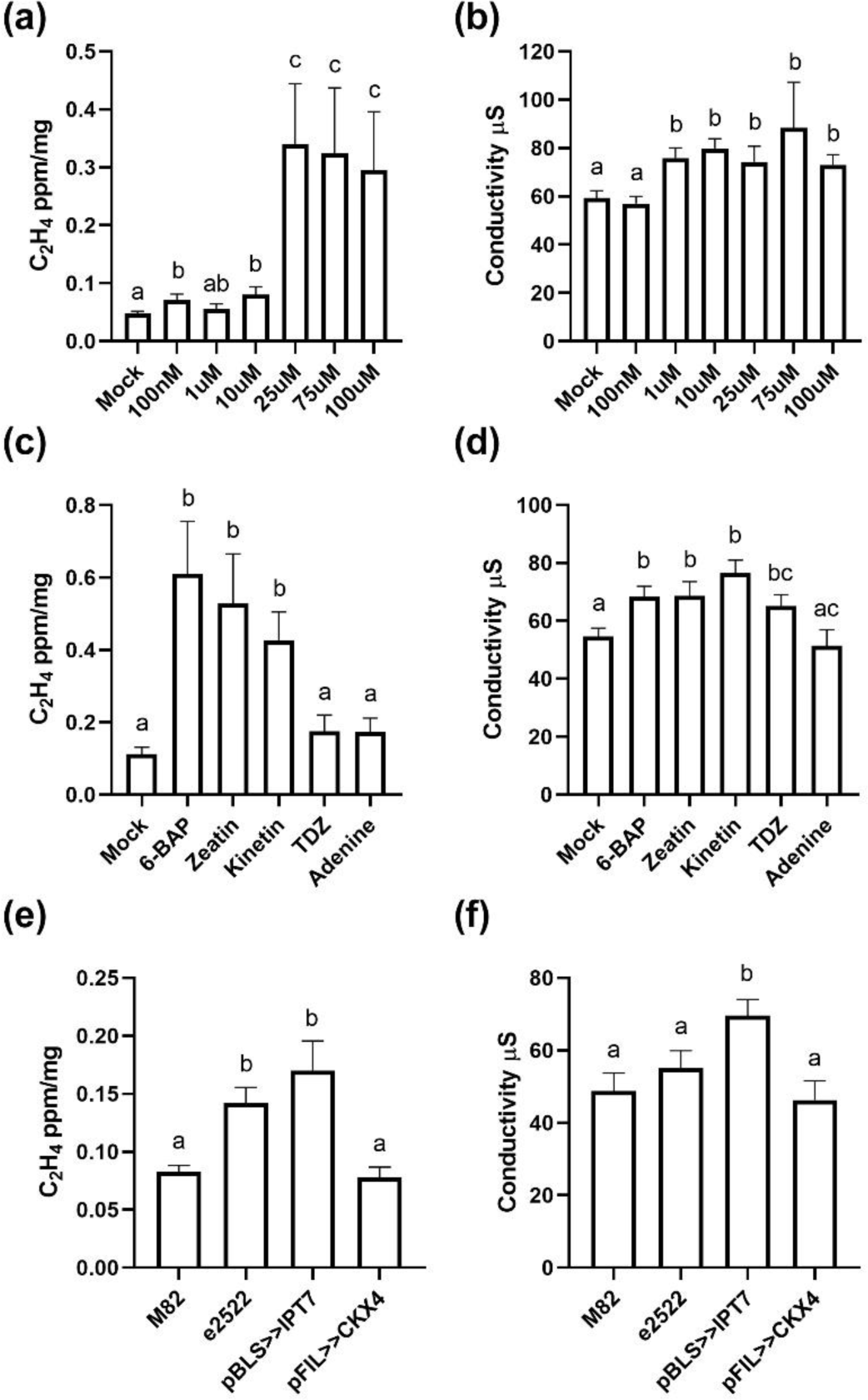
Cytokinin induces immune responses in tomato. (a)-(b) *S.lycopersicum cv* M82 leaves were treated with indicated concentrations of 6-Benzylaminopurine (6-BAP) dissolved in 1uM NaOH. (c)-(d) *S.lycopersicum cv* M82 leaves were treated with 100uM of indicated CK compounds or the control adenine. (e)-(f) Leaves of the increased CK transgene *pBLS>>IPT7*, the increased CK-sensitivity mutant *clausa e2522*, and the reduced CK content transgenic line *pFIL>>CKX4*, all in a *S.lycopersicum cv* M82 background, were assayed untreated. All assays were conducted on leaf discs as detailed in the materials section. (a,c,e) Ethylene production was measured using gas-chromatography. Average± SEM of 5 independent experiments is presented, N≥8. Different letters represent statistical significance in a two tailed t-test, p<0.039. (b,d,f) Conductivity levels of samples immersed in water for 40 h was measured. Average ± SEM of 4 independent experiments is presented, N≥7. Letters represent statistical significance in a two tailed t-test, p<0.044.

**Figure 6.**
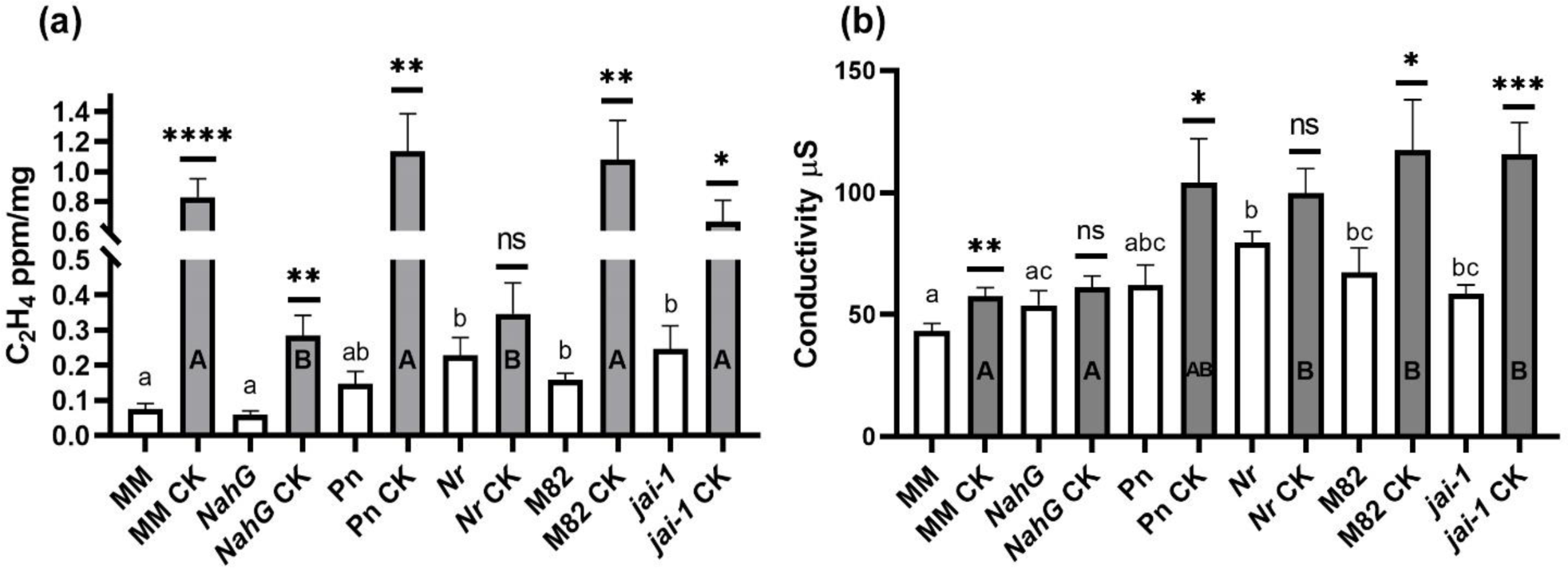
Cytokinin induced immunity is altered in salicylic acid (SA) deficient, ethylene (ET) insensitive and jasmonic acid (JA) insensitive mutants. Leaves of the SA deficient line *NahG* and its background line Moneymaker (MM), the Ethylene insensitive line Never-Ripe (*Nr*) and its background line Pearson (Pn), and the Jasmonate insensitive mutant *jai-1* and its background M82, were treated with 100uM 6BAP. (a) Ethylene production was measured using gas-chromatography. Average ± SEM of 4 independent experiments is presented, N≥8. Results were analyzed for significance in one-way ANOVA with a Tukey post hoc test, p<0.0062. Asterisks represent significance for the effect of CK within a genotype analyzed using a two-tailed t-test, *p<0.05, **p<0.01, ****p<0.0001. Lower case letters represent significant differences in baseline ethylene levels in the different genotypes, p<0.032. Upper case letters represent significant differences in ethylene production in response to CK between genotypes, p<0.0002. (b) Conductivity levels of samples immersed in water for 40 h was measured. Average ± SEM of 3 independent experiments is presented, N≥7. Results were analyzed for statistical significance in one-way ANOVA with a Dunnett post hoc test, p<0.0001. Asterisks represent significance for the effect of CK within a genotype analyzed using a two-tailed t-test, *p<0.05, **p<0.01, ****p<0.0001. Lower case letters represent significant differences in baseline conductivity levels in the different genotypes, p<0.043. Upper case letters represent significant differences in ethylene production in response to CK, p<0.037.

To examine if CK can augment defense responses elicited by a known elicitor of plant defense, we employed Ethylene Inducing Xylanase (EIX), that induces ETI in responsive cultivars (Sharon et al., 1993; Leibman-Markus et al., 2017a; Ron et al., 2000; Elbaz et al., 2002; Bar and Avni, 2009). The combination of CK and EIX induces immunity at greater levels than EIX or CK alone (see also supplemental Figure 1). We observed significant increases in ethylene production above 25uM of 6BAP added to EIX (Figure 7a). Ion leakage and ROS are also significantly increased with the addition of 6BAP when compared with EIX or CK alone (Figure 7b, supplemental Figure 2a,b). Interestingly, CK-regulated ROS homeostasis has been suggested as a possible mechanism underlying CK activated defense (Albrecht and Argueso, 2017). Kinetin also has a similar enhancing effect on EIX-induced ethylene (Figure 7c, supplemental Figure 1), while all tested CKs effect ion leakage (Figure 7d). Genotypes with elevated CK sensitivity or levels have elevated defense responses: the *IPT* overexpressor produced more ethylene in response to EIX, while both *clausa* and *IPT* had increased ion leakage and ROS production in response to EIX when compared to the background M82 cultivar (Figure 7e,f, Supplemental Figure 2c,d). The *CKX* overexpressor produced less ethylene in response to EIX.

**Figure 7.**
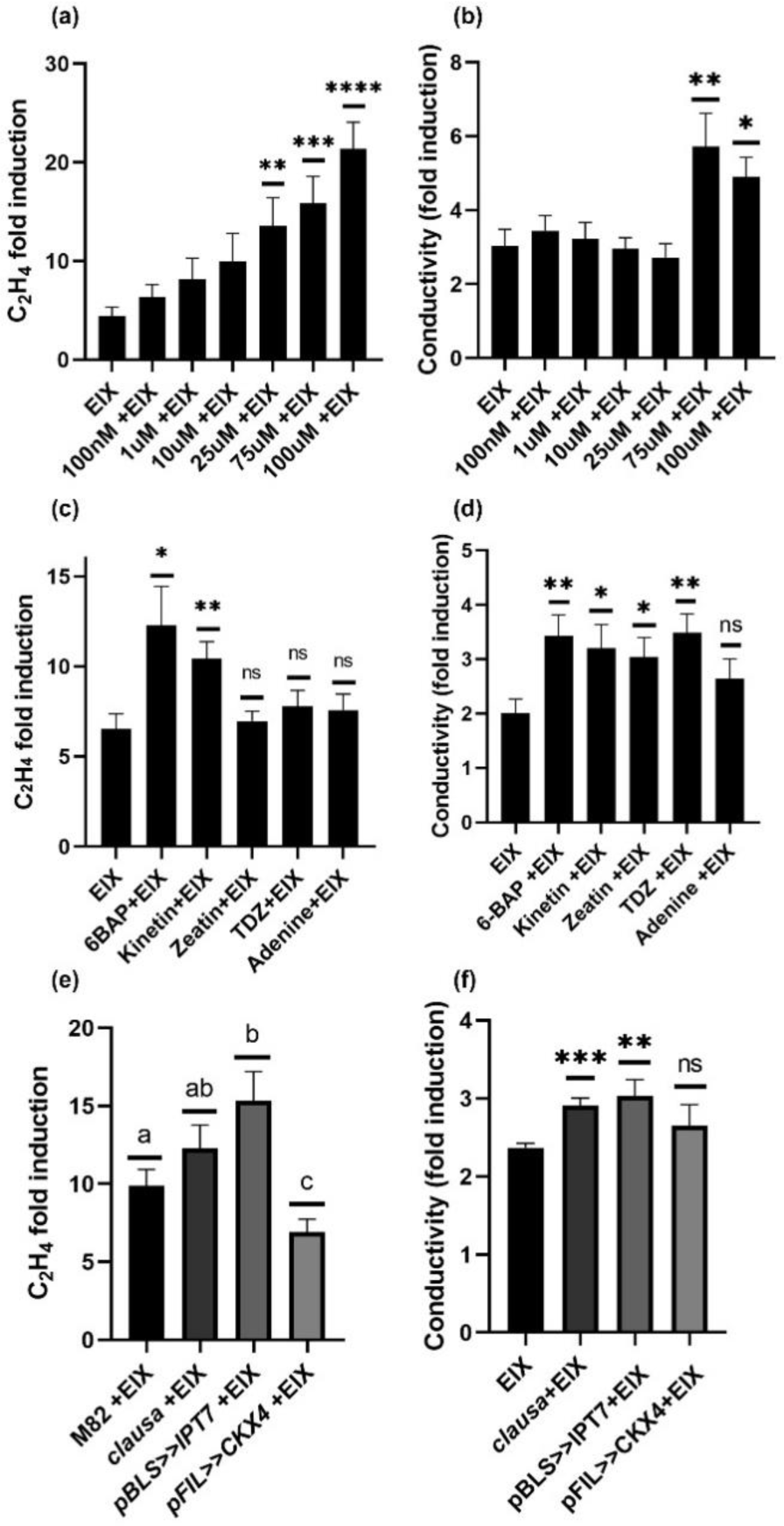
Cytokinin and Ethylene Inducing Xylanase (EIX) induce immune responses through separate pathways. (a)-(b) *S. lycopersicum cv* M82 leaves were treated with indicated concentrations of 6-Benzylaminopurine (6-BAP) dissolved in 1uM NaOH and 1 μg /mL EIX. (c)-(d) *S. lycopersicum cv* M82 leaves were treated with 100uM of indicated cytokinin (CK) compounds or the control adenine, and 1 μg/mL EIX. (e)-(f) Leaves of the increased CK transgene *pBLS>>IPT7*, the increased CK-sensitivity mutant *clausa e2522*, and the reduced CK content transgene *pFIL>>CKX4*, all in a *S. lycopersicum cv* M82 background, were treated with 1 μg/mL EIX. (a,c,e) Ethylene production was measured using gas-chromatography. Presented values are normalized to M82 mock average. Average± SEM of 5 independent experiments is presented, N≥10. Asterisks represent statistical significance in a two tailed t-test (*p <0.05, **p<0.01, ***p<0.001, ****p<0.0001). (b,d,f) Conductivity levels of samples immersed in water for 40 h was measured. Average ± SEM of 4 independent experiments is presented, N≥8. Asterisks represent statistical significance in a two tailed t-test (*p<0.05, **p<0.01, ***p<0.001).

To analyze the alterations to tomato gene expression in CK induced immunity, we examined the expression of several known defense genes in response to CK treatment, with and without subsequent pathogen inoculation. CK induces the expression of Pto-interacting 5 (*Pti-5,* Solyc02g077370), pathogenesis-related proteins (*PR1a*, Solyc01g106620) and *PR-1b* (Solyc00g174340), and pathogen induced 1 (*PI-1*, Solyc01g097270), and reduces the expression of proteinase inhibitor 2 (*PI-2*, Solyc03g020080) (Figure 8a). The addition of CK prior to *B. cinerea* inoculation causes a decrease in defense gene expression, correlating with reduced disease (Figure 8b). Positive correlations between *B. cinerea* disease levels and defense gene expression were reported previously (Harel et al., 2014). The chosen genes are all hallmarks of *B*. *cinerea* response. PI-2 and PI-1 are JA responsive and considered ISR (Martínez-Medina et al., 2013; Ament et al., 2004; Iberkleid et al., 2014; Cui et al., 2019). Pti5 is ethylene responsive, though it was found not to require ET,JA or SA for its defensive upregulation (Thara et al., 1999). PR1a is SA responsive and considered SAR (Martínez-Medina et al., 2013; López-Ráez et al., 2010). PR1b is upregulated by both SAR and ISR activation (Li et al., 2017; Harel et al., 2014). Distinctions between ISR and SAR are not clear cut in tomato, and they can overlap (Liu et al., 2016; Betsuyaku et al., 2018).

**Figure 8.**
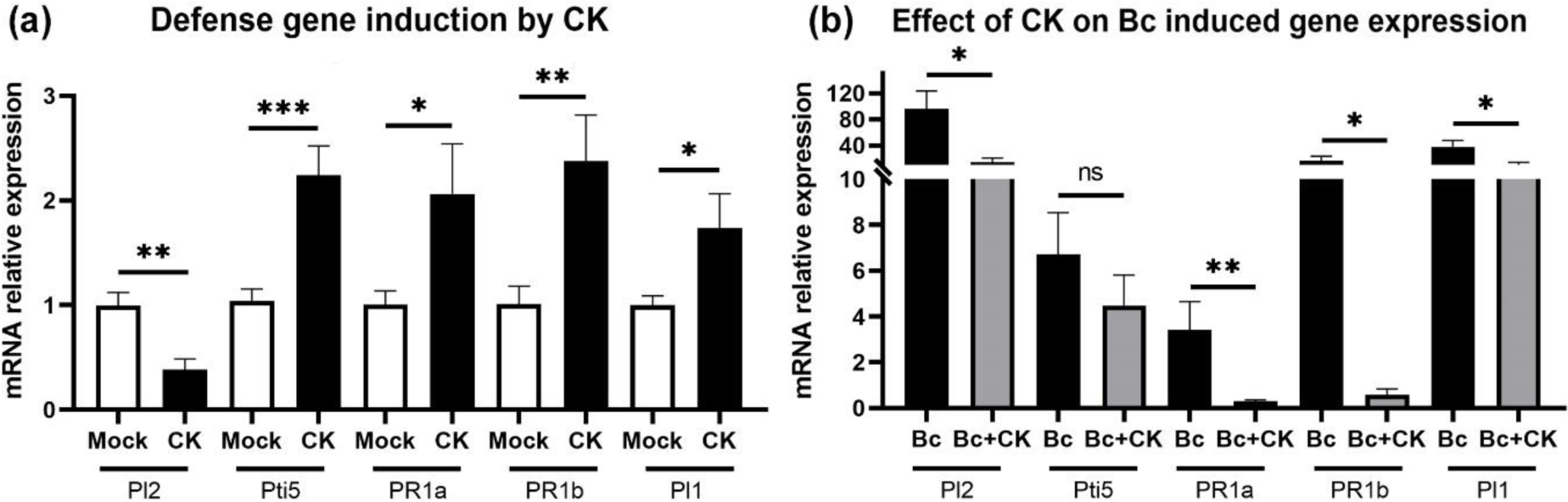
Cytokinin induces defense gene expression as a stand alone treatment and reduces defense gene expression following *B. cinerea* infection. (a) Gene expression analysis of defense genes in M82 mock and cytokinin (100uM 6-BAP) treated plants was measured by qRT-PCR. Relative expression normalized to mock. Plants treated with 1 uM NaOH were used as mock. The expression of *RPL8* was used as an internal control. Average± SEM of 4 independent experiments is shown, N≥9. (b) Gene expression analysis of defense genes in mock and CK (100uM 6BAP) treated *B. cinerea (Bc)* infected plants was measured by qRT-PCR in samples harvested 24 hours after *Bc* inoculation. Plants treated with 1 uM NaOH were used as mock. The expression of *RPL8* was used as an internal control. Relative expression normalized to untreated mock. Average ± SEM of 4 independent experiments is shown, N≥10. Results for (a) and (b) were analyzed for statistical significance in one-way ANOVA with a Dunnett post hoc test, p<0.0001 in both cases. Asterisks represent statistical significance in a two-tailed t-test comparing each gene (*p<0.05; **p<0.01; ***p<0.001).

Examining the expression of defense genes in CK altered genotypes revealed that CK affects defense gene expression endogenously as well, *Pti5, PR1b* and *PI1* being increased in with CK treatment and upon *IPT* overexpression, while *PI2* and *PR1a* are increased upon *CKX* overexpression (Figure 9a-e). Upon *B. cinerea* inoculation, *PI2*, known to correlate with *Bc* disease levels (Harel et al., 2014), is reduced in *IPT* and *clausa* (Figure 9f), while PR1b is increased (Figure 9j). With the exception of *PR1a* (Figure 9h), *clausa* behaves like *IPT* following *Bc* inoculation, similarly to its disease resistant phenotype (Figure 2).

**Figure 9.**
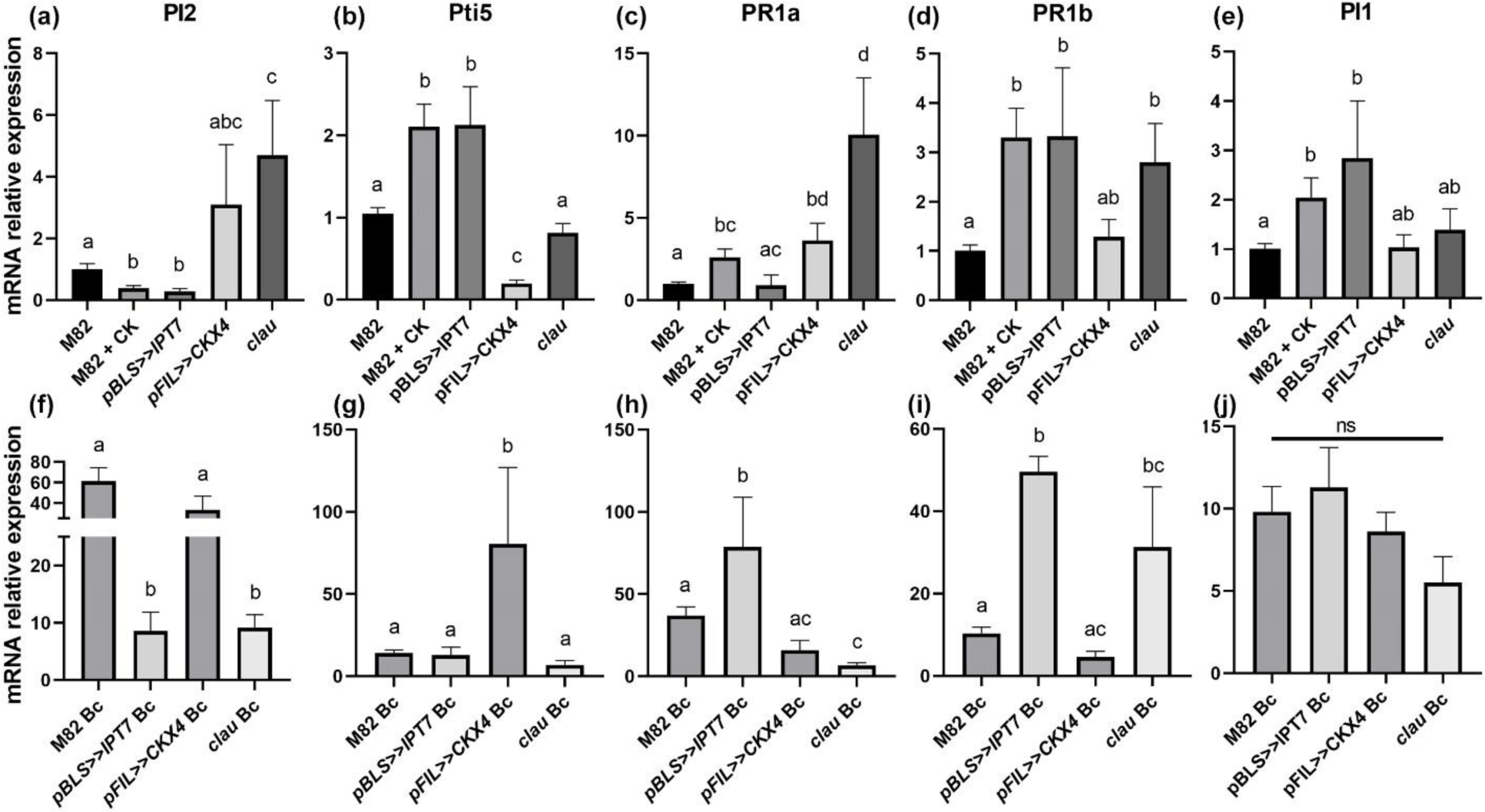
Defense gene expression in altered cytokinin genotypes in steady-state and following *B. cinerea* infection. Gene expression analysis of defense genes in the increased CK transgenic line *pBLS>>IPT7*, the increased CK-sensitivity mutant *clausa e2522*, and the reduced CK content transgenic line *pFIL>>CKX4*, as well as in M82 treated with 100uM 6BAP, in steady state (a-e) and following *B. cinerea* (Bc) infection in samples harvested 24 hours after pathogen inoculation (f-j), was measured by qRT-PCR. (a,f) proteinase inhibitor 2 (*PI2)*; (b,g) Pto-interacting 5 (*Pti5*); (c,h) pathogenesis-related proteins (*PR1a*); (d,i) *PR1b*; (e,j) pathogen induced 1 (*PI1*) genes. All samples normalized to M82 levels in steady-state. Average ± SEM of 3-5 independent experiments is shown, N≥6. The expression of *RPL8* was used as an internal control. Different letters represent statistical significance in a two-tailed t-test comparing each gene among all genotypes (p≤0.04).

### Pathogenic processes activate the CK pathway in tomato

We demonstrated that pre-treatment with CK and increased endogenous/signaling CK genotypes results in disease resistance in tomato (Figures 1-2). Is this an endogenously employed mechanism in tomato? Do tomato plants activate their CK machinery upon pathogen attack? To answer this question, we examined endogenous modulations to the CK pathway during pathogenesis.

During *B. cinerea* infection, the expression of CK responsive type-A Tomato Response Regulators (TRRs) increases and the expression of *CKX* genes is also significantly altered (Figure 10a). 48 hours after Bc inoculation, the amount of active CKs decreases, significantly in the case of trans-Zeatin and iso-Pentenyl-Riboside (Figure 10b). Using the CK response marker TCS (Zürcher et al., 2013; Bar et al., 2016), we determined that the CK pathway is activated upon *Bc* infection in mature leaf tissue, peaking at 48 hours after inoculation (Figure 10c-d). *B. cinerea* droplet inoculation causes a specific response in the leaf tissue in contact with the fungus; following the spread of infection and necrosis of the tissues, the CK responsive halo spreads out from the cite of pathogen inoculation (Figure 10d). Mock treated leaves (droplet “inoculation” with infection media) produced little to no TCS signal (Figure 10c-d).

**Figure 10.**
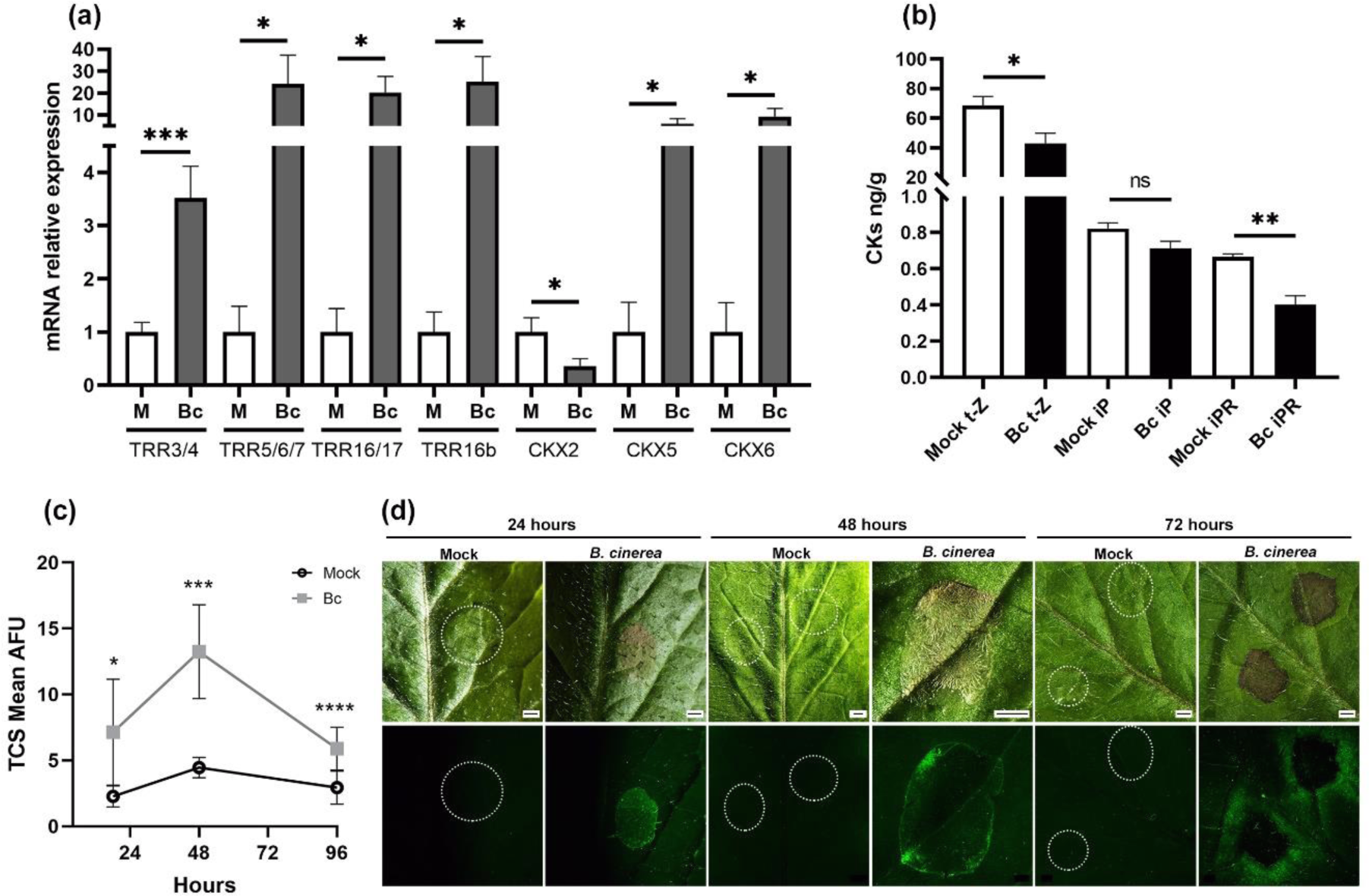
Pathogen infection induces host cytokinin response. (a) Gene expression analysis of cytokinin (CK) responsive genes in M82 mock (M) and *B. cinerea* (*Bc)* infected samples 24h after inoculation was measured by qRT-PCR. Plants treated with 1 uM NaOH were used as mock. Relative expression normalized to mock. Average ± SEM of 3 independent experiments is shown, N=6-15. Asterisks represent statistical significance in a two-tailed t-test (*p<0.05; ***p<0.001). (b) LC/MS quantification of the active CKs trans-zeatin (t-Z), iso-pentenyl (iP) and iso-pentenyl-riboside (iPR) 24 hours after *Bc* inoculation. Plants treated with 1 uM NaOH were used as mock. Average ± SEM is shown, N=3-6. Asterisks represent statistical significance in a one-way ANOVA test with a Bonferroni post hoc test, p<0.0001 (*p<0.05, **p<0.01). (c-d) Stereomicroscope analysis of *pTCS::3XVENUS* expression surrounding *Bc* inoculation site at indicated time points post droplet inoculation. Plants treated with 1 uM NaOH were used as mock. (c) Fluorescence quantified using ImageJ, average± SEM of 3 independent experiments is shown, N=4-15. Asterisks represent statistical significance in a two-tailed t-test (*p<0.01; ***p<0.001, ****p<0.0001). (d) Representative images shown, bar=250um.

### CK-regulated PRR presence on endomembrane compartments mediates CK-induced disease resistance

How does CK affect immune signalling and disease resistance? Other than the evidence provided here (Figure 3) and by others (Argueso et al., 2012; Choi et al., 2010; Naseem et al., 2012) that SA is required for CK induced immunity, partially through binding of the CK responsive ARR2 to the SA pathway activator TGA3 (Choi et al., 2010), and that phytoalexins can play a part (Grosskinsky et al., 2011)-no additional cellular mechanisms have been reported. Pattern Recognition Receptors, PRRs are the first line of defence and immune-activation in plant cells. Based on the evidence in the literature that CK modulates endocytic trafficking of PIN1 in its regulation of auxin (Marhavý et al., 2011), we were prompted to investigate whether CK may also affect trafficking of immune receptors as a possible mechanism for promoting immune responses and disease resistance. Since we determined that CK enhances EIX-induced immune responses (Figures 7, S1, S2), we examined whether it affects trafficking of the PRR LeEIX2, the EIX receptor (Ron and Avni; Bar and Avni, 2009). As can be seen in Figure 11, CK enhances both the endosomal presence (Figure 11a,k) and vesicular size (Figure 11b,k) of LeEIX2 endosomes, without affecting total cellular content of the protein (Figure 11c). Upon EIX treatment, which enhances endosomal presence (Figure 11d,h) and size (Figure 11e,h) of LeEIX2 in the mock treated samples, as previously reported (Pizarro et al., 2018), there is no further increase with the addition of CK (Figure 11d,e,l), though the level of the receptor in the cell appears to increase slightly with the combination of both treatments (Figure 11f). CK enhances cellular immunity (Figures 5, 8), and LeEIX2 presence on endosomes (Figure 11 a-l). LeEIX2 endosomal presence is required for EIX induced immunity (Ron and Avni; Bar and Avni, 2013; Sharfman et al., 2011), which CK enhances (Figure 7). Does this CK-mediated enhancement of PRR endosomal presence act as a mechanism for increased disease resistance? To examine this, we used a SlPRA1A overexpressing line, which was previously shown to have a decreased presence of Receptor Like Protein (RLP)-type PRRs in the cell plasma membrane, due to receptor degradation, along with a reduction in LeEIX2-mediated immune responses (Pizarro et al., 2018). Wild type (WT), 35S driven GFP overexpressing, and 35S driven SlPRA1A-GFP overexpressing *Solanum lycopersicum cv* M82 tomato plants were treated with 100uM 6-Benzyl Amino Purine and infected with B.cinerea 24 hours later. Disease was assessed 7 days after inoculation. CK pre-treatment significantly decreased *B. cinerea* disease levels in WT M82 plants (Figure 11m; see also Figures 1-2), as well as in plants overexpressing GFP. However, plants overexpressing SlPRA1A, which have decreased levels of RLP type PRRs, are not responsive to CK, and no reduction in *B. cinerea* disease in these plants is observed upon CK treatment (Figure 1m).

**Figure 11.**
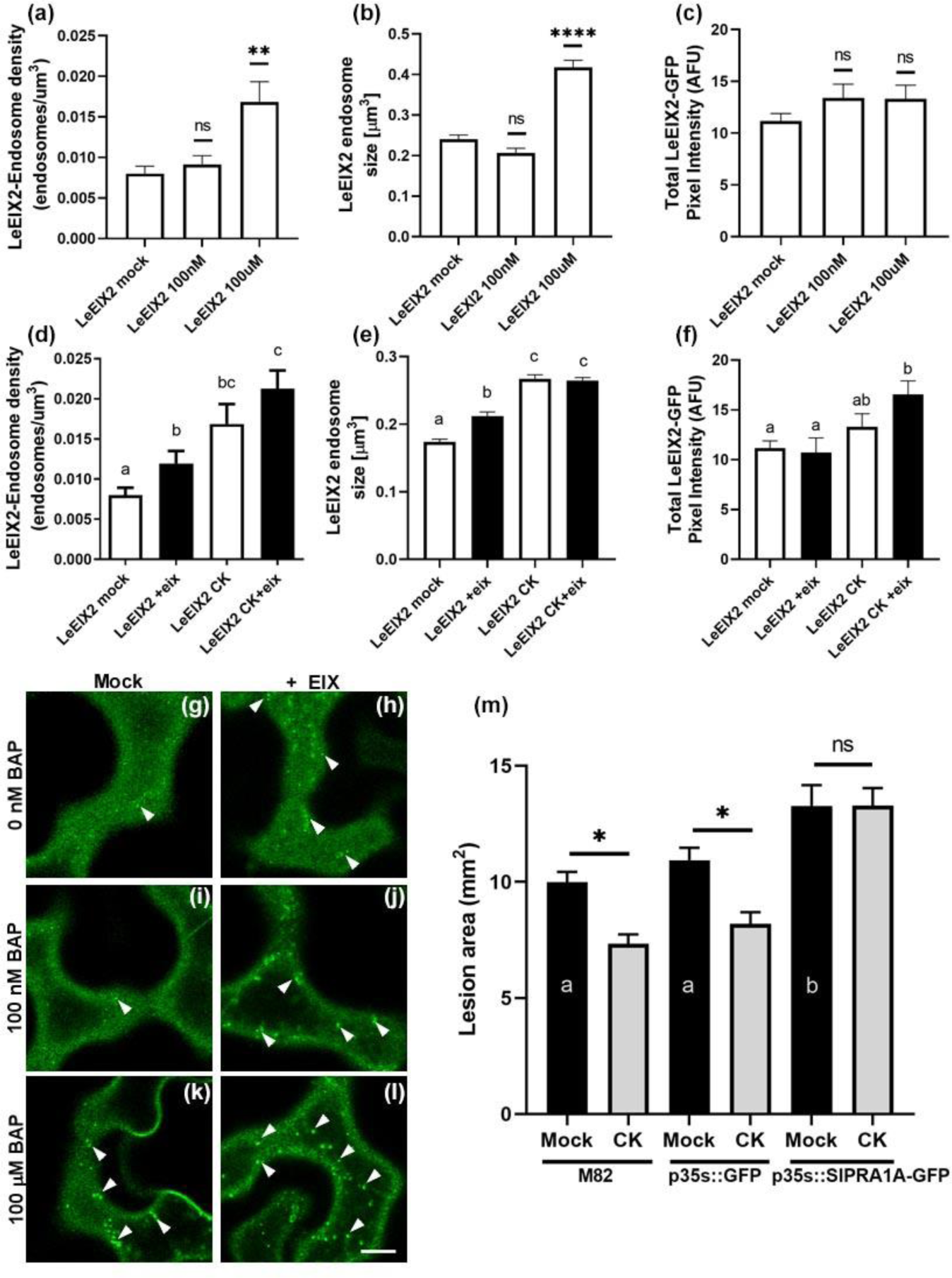
Cytokinin enhances cellular trafficking of the PRR LeEIX2. *N. benthamiana* epidermal cells transiently expressing LeEIX2-GFP were treated with CK or Mock as indicated (a-l), and subsequently treated with 1mg/mL EIX (d-f,h,j,l) for 5 minutes, followed by live cell imaging. (a-f) Graphs depicting the analysis of confocal microscope images acquired using a Zeiss LSM780 confocal microscope system with a C-Apochromat 40×/1.2 W Corr M27 Objective, using a 488 nm excitation laser (3% power, 493–535 nm emission range). (g-l) Representative images taken from the membrane/endosomal plane, bar=10um. (a-c,g-k) Effect of CK (6BAP, concentrations as indicated) on endosomal presence (a,g-k) vesicular size (b,g-k), and total GFP cellular content of LeEIX2 (c). (d-f,h-l) Effect of the combination of CK and EIX (1mg/mL) on endosomal presence (d,h-l), vesicular size (e,h-l) and total GFP cellular content of LeEIX2 (f). Image analysis (21-38 images per treatment) was conducted with Fiji-ImageJ using the raw images and the 3D object counter tool for quantifying endosome numbers and size, and the measurement analysis tool for quantifying pixel intensity. Graphs represent average ±SEM, N>20 per treatment. Statistical significance was determined in a one-way ANOVA with a Dunnett post hoc test, p=0.0005 (a), p<0.0001 (b,d,e), p=0.26 (c), p=0.056 (f). Asterisks (a-c) represent statistically significant differences from the mock treatment (**p<0.01, ****p<0.0001). Letters (d-f) represent statistically significant differences between samples (p≤0.038 where different letters are indicated). (m) *S.lycopersicum cv* M82 plants, WT or expressing GFP or SlPRA1A, both driven by the 35S promoter, were spray-treated with 100uM 6-Benzylaminopurine (6-BAP) dissolved in 1uM NaOH, and inoculated with 10 ul of *B. cinerea* spore solution (10^6^ spores ml^-1^) 24 hours later. Lesion area was measured 7 days after *B. cinerea* inoculation using ImageJ. Graph represents the results of 4 independent experiments ±SEM, N≥20 for each genotype/ treatment combination. Results were analyzed for statistical significance using one-way ANOVA with a Bonferroni post hoc test, p<0.0001. Asterisks indicate statistically significant differences between Mock and CK treatment, letters indicate statistically significant differences in a two-tailed t-test conducted among mock samples only.

## Discussion

Several previous reports have indicated that endogenous and exogenous CK treatment can result in resistance to pathogens (reviewed in (Albrecht and Argueso, 2017). Our work supports the notion that the main mode of action for CK induced pathogen resistance is through induced immunity (Figures 2, 5). In Arabidopsis, cytokinin-treated plants demonstrated upregulation of defense gene expression and callose deposition coupled with decreased pathogen growth (Choi et al., 2010; Argueso et al., 2012). CK treatment alone was also previously shown to induce ethylene biosynthesis (Coenen and Lomax, 1998) and *PR1a* expression (Choi et al., 2010).

Here, we show that CK acts systemically to induce immunity to foliar fungal pathogens when applied by soil drench to tomato roots (Figure 2f). CK induces ethylene biosynthesis and ion leakage, both exogenously, using different CK compounds, and endogenously, using CK mutants. CK also induces defense gene expression in tomato (Figures 8-9), as was previously reported for Arabidopsis (Choi et al., 2010; Argueso et al., 2012) and rice (Jiang et al., 2013). Interestingly, the MYB transcription factor mutant *clausa*, that has increased CK sensitivity and behaves like an overexpression of *IPT* developmentally (Bar et al., 2016), possesses decreased amounts of endogenous CK. Despite decreased endogenous CK, *clausa* retains the CK protective effect and exhibits increased defense responses (Figure 5) and pathogen resistance (Figure 2), indicating that CK induced immunity is dependent on host signaling pathways. Although variable in steady state, upon *B. cinerea* infection, the defense gene expression pattern in *clausa* resembles that of the *IPT* overexpressing line, in agreement with increased pathogen resistance observed in both genotypes. This demonstrates that defense gene expression in steady state can be uncoupled from subsequent pathogen resistance in genetically primed plants in certain cases, and testifies to the flexibility of host responses.

Previous results coupled with our work suggest that cytokinin can be viewed as a priming agent, acting to potentiate defense responses (Choi et al., 2010; Argueso et al., 2012). Consistent with the notion that priming agents have a low level, often transient effect on host defense physiology (Conrath et al., 2015), CK induces much lower levels of defense gene expression than *B. cinerea* (compare Figure 8a to Figure 8b). Interestingly, comparing CK with EIX, a well-known elicitor of plant defense responses, a priming agent, demonstrates that ≥ 25uM CK induces ethylene biosynthesis to similar levels as EIX, confirming that the plant responds to CK as it would to a priming agent.

Though mechanisms of action were reported to differ in different host plants (Choi et al., 2010; Grosskinsky et al., 2011; Albrecht and Argueso, 2017), it is generally accepted that, in Arabidopsis, CK induces immunity to biotrophic pathogens through SA dependent pathways. In agreement, our results demonstrate that a functioning SA pathway is required to achieve CK induced pathogen resistance (Figure 3b,h,m). CK was able to induce ethylene biosynthesis, to a lesser level than in the WT, in a SA deficient background (Figure 6a), demonstrating that *NahG* plants have a low level of CK priming that is significant but insufficient to promote disease resistance. Interestingly, though we have shown that CK induced pathogen resistance requires the SA pathway (Figure 3) absolute SA levels or *Bc*-mediated SA content reduction do not directly correspond with pathogen resistance (Figure 4), suggesting that CK-mediated pathogen resistance may require additional signaling mechanisms.

We also show here that normal ethylene sensitivity is required for CK induced immunity and disease resistance in tomato (Figures 3, 6). Defense responses induced by CK (Figure 6) were compromised in an ethylene sensitivity deficient background. Finally, we found that JA sensitivity is not required for CK induced pathogen resistance (Figures 3, 6).

Distinctions between SAR and ISR are not always clear-cut. Occurrences of overlaps and/or co-activation between these pathways have been previously reported (Liu et al., 2016; Betsuyaku et al., 2018). Our work suggests that in tomato, CK activates systemic resistance that requires SA signaling (Ryals et al., 1996). This is supported by the fact that CK induced immunity requires a functioning SA pathway. Further support comes from the fact that CK and EIX, an elicitor protein derived from the JA pathway ISR elicitor Trichoderma, (Shoresh et al., 2005), augment the levels of defense elicited by each alone (supplemental Figure 1), indicating that they potentiate plant immunity through separate pathways. However, evidence of overlap between SAR and ISR also exists in the context of CK, with CK activating classical ISR genes (Figure 8a), and requiring ET, though not JA, to induce pathogen resistance (Figure 3). This is perhaps not surprising, given that CK signaling was reported to be upstream of ethylene in several cases (Zdarska et al., 2015; Robert-Seilaniantz et al., 2011).

Our work demonstrates that CK affects internalization of the PRR LeEIX2, which mediates immune responses to the xylanase EIX (Ron and Avni). LeEIX2 was shown to be internalized following ligand application (Ron and Avni), and to require this internalization for a full mounting of defense responses (Bar and Avni, 2009). Interestingly, CK increased both LeEIX2 internalization and EIX mediated defense responses (Figures 7, 11, S1, S2), suggesting that increased internalization could be at least one of the mechanisms underlying the increase in immune responses. Supporting this hypothesis, we found that in the SlPRA1A overexpressing line, in which expression and plasma membrane presence of RLP-type PRRs, including LeEIX2, are greatly reduced (Pizarro et al., 2018), CK is no longer able to mediate resistance to *B. cinerea*, suggesting that it requires normal levels of PRRs to do so.

Cytokinin-based direct regulation of receptor endocytosis has been shown for PIN1 (Marhavý et al., 2011), where the authors determined that this endocytic regulation is a specific mechanism to rapidly modulate the auxin distribution in cytokinin-mediated developmental processes, through a branch of the cytokinin signaling pathway that does not involve transcriptional regulation. Therefore, in addition to regulating immunity and diseases resistance through the SA pathway, CK may regulate endocytic trafficking independent of transcriptional regulation, accounting for the rapid plant response.

To the best of our knowledge, our work is the first one reporting that CK induces resistance to *B. cinerea* and *O. neolycopersici* in tomato. This also points to an overlap between SAR and ISR in the case of CK induced immunity, as the host plant requires JA signaling to resist necrotrophic pathogens such as *B. cinerea* (Thomma et al., 1998; Durrant and Dong, 2004; Liu et al., 2016) (see also Figure 3f,m), yet CK is able to prime this resistance. Our work contradicts results achieved in tobacco (Grosskinsky et al., 2011), a solanaceous host, in two respects: first, SA signaling was found not to be required for CK induced immunity to *Pst* and, second, CK was found not to induce resistance to *B. cinerea*. In that work, the authors found significant roles for the phytoalexins scopoletin and capsidiol in CK induced immunity to *Pst* in tobacco; the lack of SA pathway requirement in that particular system, could be attributed to the time-course of infection and phytolaexin production (Albrecht and Argueso, 2017). The lack of protectant effect for CK against *B. cinerea* in tobacco may stem from different host biology, though more likely, it originates in differences in experimental design.

Why does CK induce immunity in plants? Are classical developmental hormones also “defense” hormones, or is CK induced immunity attributable to hormonal crosstalk? The swiftness of CK-induced immune processes seems to indicate that CK action is relatively “direct” (see Figures 5,7), and supports the idea of regulation of PRR trafficking as one of the underlying mechanisms. In developmental contexts, CK can be viewed as a “juvenility” factor, promoting meristem maintenance and morphogenetic processes, and delaying differentiation and senescence (Gordon et al., 2009; Kurakawa et al., 2007). Simplistically, could senescence-like processes activated by pathogen derived tissue destruction in both biotrophic and necrotrophic infections make it evolutionarily economical for the plant to adapt those pathways for use in the war against pathogens, by recognizing levels of self-CK as a signal to activate immunity? Certainly, delayed senescence/ enhanced juvenility can correlate with increased pathogen resistance (Pogány et al., 2004; Grosskinsky et al., 2011). It is worth nothing that, although CK signaling it is not normally active in mature, differentiated tissue such as a mature tomato leaf (Farber et al., 2016; Shani et al., 2010), upon pathogen attack, the CK system is activated (see Figure 10), suggesting that CK defensive function could be spatio-temporally regulated. CK signaling outside of a defined developmental window in specific tissues could facilitate the translation of CK into a defense signal. Thus, it would seem that the previously accepted paradigm, that pathogenesis processes cause a shunting of available plant resources towards immunity, shutting off growth programs to divert all available resources towards defense (Karasov et al., 2017; Berens et al., 2017), can be updated to reflect that some “developmental programs” are not shut off but rather, modified and appropriated for defense purposes.

## Acknowledgements

The authors would like to thank Yigal Elad for the *B.cinerea* strain, and seeds of *NahG* and *Nr* mutants; Naomi Ori for seeds of *clausa*, *pFIL>>CKX*, *pTCS::3XVENUS*, *jai-1*; Adi Avni for EIX and seeds of p35S::SlPRA1A-GFP; Tzahi Arazi for seeds of p35S::GFP; and Mira Weissberg-Carmeli and Felix Shaya for the metabolomic analyses. Research in the Bar group is supported by the Israeli ministry of agriculture. RG is supported by the Indo-China ARO Postdoctoral Fellowship Program. MB thanks members of the Bar group for continuous discussion and support. Publication No XXX/19 of the ARO.

## Author Contributions

MB and RG conceived and designed the study. RG, LP, ML-M, and IM formulated the methodology and carried out the experiments. RG, LP, ML-M, and MB analyzed the data. All authors contributed to the writing of the manuscript.

## Materials and Methods

### Plant materials and growth conditions

Seeds of the *S. lycopersicum* cultivar M82 were used throughout the study. Tomato mutant and transgenic lines, which were employed in the assays were as follows: seeds of the increased CK: *pBLS>>IPT7*, decreased CK: *pFIL>>CKX4*, increased CK sensitivity: *clausa* (e2522), and JA insensitive *jai-1*, all in an M82 background, were obtained from Prof. Naomi Ori, the Hebrew University of Jerusalem (Shani et al., 2010; Bar et al., 2016). Seeds of the decreased SA: *NahG* and its parental WT *Moneymaker*, decreased ethylene sensitivity: never ripe (*Nr*) and its parental WT *Pearson* were obtained from Prof. Yigal Elad (Mehari et al., 2015). Plants were grown from seeds in soil (Green Mix; Even-Ari, Ashdod, Israel) in a growth chamber, under long day conditions (16 hr:8 hr, light:dark) at 24°C.

### Cytokinins treatments

Cytokinin (CK) (6-benzyl aminopurine: BAP) (Sigma-Aldrich) was sprayed onto 4-5 week-old plants, or soil drenched onto the roots (100ml 15cm^-1^ diameter pot). BAP solutions were prepared from a stock in 1uM NaOH, and diluted into an aqueous solution to the desired BA concentration, with the addition of Tween 20 (100ul l^-1^). Mock plants were sprayed or soil-drenched with the aforementioned solution of NaOH with Tween20. The CK analogs kinetin (6-Furfurylaminopurine riboside), trans-zeatin [6-(4-hydroxy-3-methylbut-2-enylamino) purine] and thidiazuron (TDZ), as well as the control adenine (all from Sigma-Aldrich) were prepared in 1uM NaOH (kinetin and zeatin), 1M HCl (adenine) and 100 % dimethyl sulfoxide (thidiazuron). Pathogen inoculations were carried our 24 hours after spray treatments and 3 days after soil drench, as detailed above.

### Pathogen infection and disease monitoring

Pathogenesis assays were conducted both on whole plants (Figure 1, 2g, 4, 8, 9, 10) and on detached leaves (Figure 1g, 2, 3, 11). For disease assays conducted on whole plants, inoculation was conducted as described below, and, as indicated, leaflets were photographed on the plant 5-10 days post inoculation (Figure 1f-h, 2f), or removed and immediately photographed (Figures 1a-e, 10c-d). For gene expression analyses (Figures 8, 9, 10a), leaf tissue (1 cm diameter around the inoculation site) was removed 24 hours after inoculation and immediately processed for RNA/ cDNA preparation and qRT-PCR analyses (see below). For metabolomic analyses (Figures 4, 10b), leaf tissue (entire spray-inoculated leaflets) was removed 48 hours after inoculation and immediately processed for hormone extraction (see below).

******B. cinerea****** (isolate BcI16) cultures were maintained on potato dextrose agar (PDA) (Difco Lab) plates and incubated at 22 °C for 5-7 days. *B. cinerea* spores were harvested in 1 mg mL^-1^glucose and 1 mg mL^-1^ K2HPO4 and filtered through cheesecloth. Spore concentration was adjusted to 10^6^ spores ml^-1^ using a hemocytometer. Each tomato leaflet was either spray inoculated with the spore suspension, or inoculated with two droplets of 10 μl spore suspension-as indicated.

******O. neolycopersici****** was isolated from young leaves of tomato plants grown in a commercial greenhouse. Conidia of *O. neolycopersici* were collected by rinsing infected leaves with sterile water. Concentrations of these conidial suspensions were determined under a light microscope using a hemocytometer. The conidia suspensions were adjusted to 10^4^ ml^-1^ and then sprayed onto plants at 5 mL per plant. All suspensions were sprayed within 10 to 15 minutes of the initial conidia collection. Suspensions were applied with a hand-held spray bottle and plants were left to dry in an open greenhouse for up to 30 minutes.

Inoculated plants were kept in a temperature controlled growth chamber at 22°C, and inoculated excised leaves were kept in a humid growth chamber at 22°C. Controls consisted of plants or leaves treated with water/buffer without pathogen inoculation. The area of the necrotic lesions or % of infected leaf tissue was measured five to ten days post inoculation using ImageJ.

### Plant immunity assays

Immunity assays (Figures 5-7, S1, S2) were conducted on leaf discs from indicated genotypes.

## Ethylene measurement

Ethylene production was measured as previously described (Leibman-Markus et al., 2017a). Leaf discs 0.9 cm in diameter were harvested from indicated genotypes, and average weight was measured for each plant. Discs were washed in water for 1-2 h for EIX and steady state assays or incubated for 3-4 h in different concentration of CK. Every six discs were sealed in a 10 mL flask containing 1 ml assay medium (with or without 1 μg ml^-1^ EIX or with or without CK) for 4 h (for EIX) or overnight (for CK) at room temperature. Ethylene production was measured by gas chromatography (Varian 3350, Varian, California, USA).

## Conductivity measurement

Leaf discs 0.9 cm in diameter were harvested from indicated genotypes. Discs were washed in a 50 ml water tube for 3 h. Every five discs were floated in a 12-well plate containing 1 ml of water, with or without 1 μg ml^-1^ EIX or with or without CK, adaxial surface down, at room temperature with agitation. Conductivity was measured in the water solution after 40 h of incubation using an conductivity meter (EUTECH instrument con510).

## Measurement of ROS generation

ROS measurement was measured as previously described (Leibman-Markus et al., 2017b). Leaf discs 0.5 cm in diameter were harvested from indicated genotypes. Discs were floated in a white 96-well plate (SPL Life Sciences, Korea) containing 250 μl distilled water for 4–6 h at room temperature. After incubation, water with and without different concentrations of CK and its analogs was removed and a ROS measurement reaction containing either 1 μg ml^-1^ EIX or water (mock) was added. Light emission was measured immediately and over indicated time using a luminometer (Turner BioSystems Veritas, California, USA).

### RNA extraction and qRT-PCR

Plant total RNA was extracted from tomato plants 24 hours after *B. cinerea* inoculation using Tri reagent (Sigma-Aldrich) according to the manufacturer’s instructions. RNA was isolated from plants infected as indicated (whole plant assays). RNA (3μg) was converted to first strand cDNA synthesis using reverse transcriptase (Promega, United States) and oligodT ^15^. qRT-PCR was performed according to the Power SYBR Green Master Mix protocol (Life Technologies, Thermo Fisher, United States), using a Rotor-Gene Q machine (Qiagen). Supplemental Table 1 lists the specific primers used in this work. Relative expression quantification was calculated using copy number method for gene expression experiments (D’haene et al., 2010). The housekeeping gene coding for ribosomal protein RPL8 (accession number Solyc10g006580) was used for the normalization of gene expression in all analyses.

### Phytohormone analysis

Hormone extraction was performed according to (Shaya et al., 2019). Plants were inoculated with *B. cinerea* as described above. 48 hours after inoculation, entire leaflets were harvested from the inoculated plants. Phytohormones were quantified in the harvested tissue. Briefly, frozen tissue was ground to a fine powder using a mortar and pestle. 200-450 mg powder was transferred to a 2ml tube containing 1ml extraction solvent (ES) mixture (79% IPA: 20% MeOH: 1% acetic acid) supplemented with 20ng of each deuterium-labelled internal standard (IS, Olomouc, Czech Republic). The tubes were incubated for 60min at 4°C with rapid shaking and centrifuged at 14,000g for 15min at 4°C. The supernatant was collected and transferred to 2mL tubes. 0.5ml of ES was added to the pellet and the extraction steps were repeated twice. The combined extracts were evaporated using speed-vac at RT. Dried samples were dissolved in 200μl 50% methanol and filtered with 0.22μm cellulose syringe filter. 5–10μL were injected for each analysis. LC–MS-MS analyses were conducted using a UPLC-TripleQuadrupoleMS (WatersXevo TQMS). Separation was performed on Waters Acuity UPLC BEH C18 1.7μm 2.1×100mm column with a VanGuard precolumn (BEH C18 1.7μm 2.1×5mm). The mobile phase consisted of water (phase A) and acetonitrile (phase B), both containing 0.1% formic acid in the gradient elution mode. The flow rate was 0.3ml/min, and the column temperature was kept at 35 °C. Acquisition of LC–MS data was performed using MassLynx V4.1 software (Waters). Quantification was done using isotope-labeled internal standards (IS). Solvent gradients and MS-MS parameters are detailed in supplemental Table 2.

### Imaging of CK–response synthetic promoter pTCS:3XVENUS

Tomato plants expressing the synthetic promoter pTCS:3XVENUS (Bar et al., 2016; Zürcher et al., 2013) were spot inoculated with *B. cinerea* and kept at 22◦C under long day conditions (16 h light/8 h dark) for 1 to 3 days. VENUS expression was analyzed using a Nikon SMZ-25 stereomicroscope equipped with a Nikon-D2 camera and NIS elements v5.11 software. ImageJ software was used for analysis and quantification of captured images.

### Trafficking imaging and analysis

*N. benthamiana* epidermal cells transiently expressing LeEIX2-GFP were infiltrated with BAP 100nM, BAP 100 µM or Mock (distilled water with 0.2% Tween). Leaf discs of 1cm diameter from the different conditions (6 hours after infiltration) were treated with EIX (1mg/mL) for 5 minutes and live cell imaging was conducted. A total of 8 plants per treatment were studied in three separate experiments. Confocal microscopy images were acquired using a Zeiss LSM780 confocal microscope system with a C-Apochromat 40×/1.2 W Corr M27 Objective. GFP images were acquired using a of 488 nm excitation laser (3% power), with the emission collected in the range of 493–535 nm. Images of 8 bits and 1024 × 1024 pixels were acquired using a pixel dwell time of 1.27, pixel averaging of 8, and pinhole of 1 airy unit. Image analysis (21-38 images per treatment) was conducted with Fiji-ImageJ using the raw images and the 3D object counter tool for quantifying endosome numbers, and the measurement analysis tool for quantifying pixel intensity (Schindelin et al., 2012).

### Statistical Analyses

All data is presented as average ±SEM. Differences between two groups were analyzed for statistical significance using a two-tailed t-test. Differences among three groups or more were analyzed for statistical significance with a one-way ANOVA. Regular ANOVA was used for groups with equal variances, and Welch’s ANOVA for groups with unequal variances. When a significant result for a group in an ANOVA was returned, significance in differences between the means of different samples in the group were assessed using a post-hoc test. Tukey was employed for samples with equal variances when the mean of each sample was compared to the mean of every other sample. Bonferroni was employed for samples with equal variances when the mean of each sample was compared to the mean of a control sample. Dunnett was employed for samples with unequal variances. All statistical analyses were conducted using Prism8^TM^.

---

**Supplemental Figure 1.**
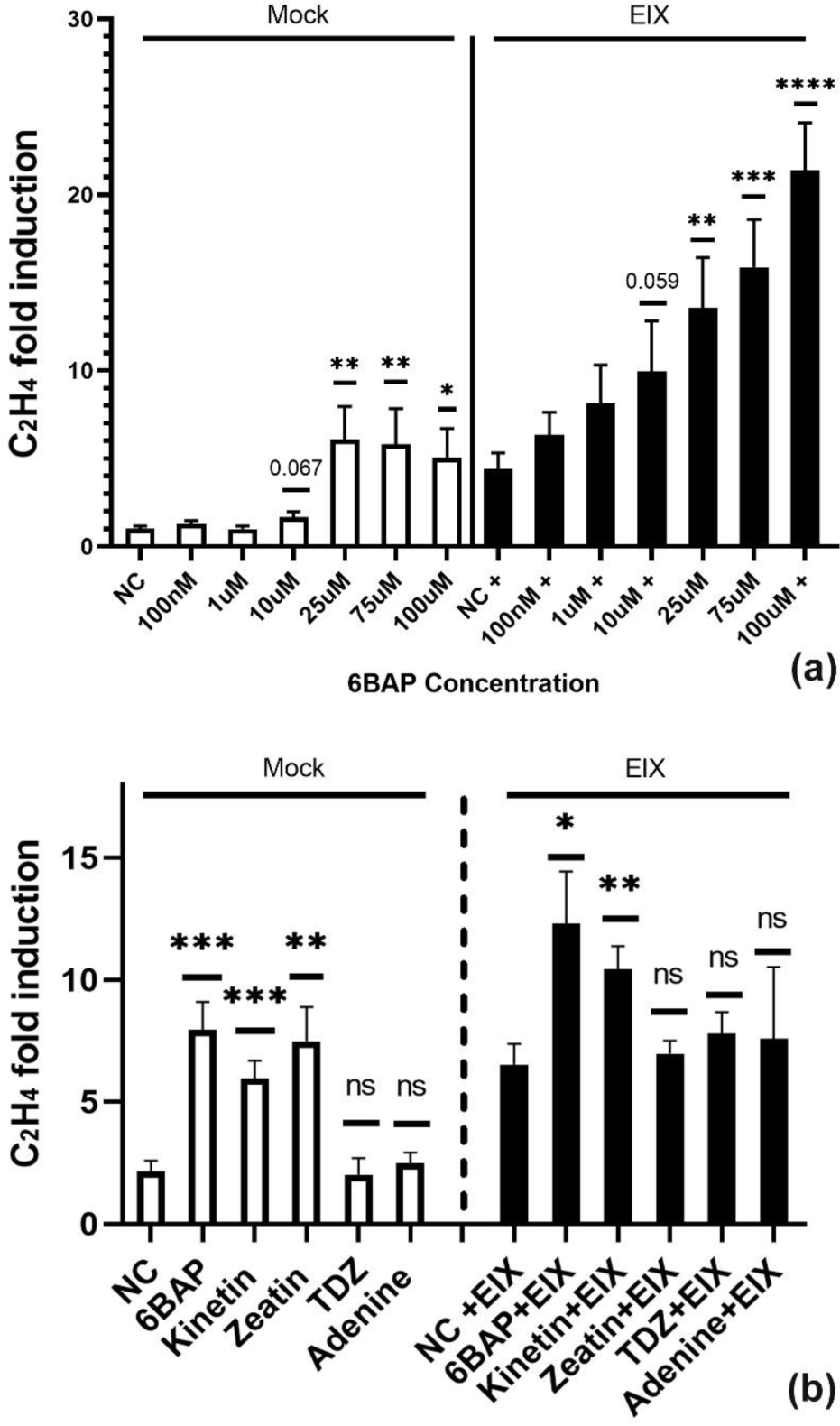
Comparison of cytokinin (CK) induced immunity and Ethylene Inducing Xylanase (EIX)+CK induced immunity. (a) *S. lycopersicum cv M82* leaves were treated with indicated concentrations of 6-Benzylaminopurine (6-BAP) or (b) with 100uM of different CK compounds or the control adenine, alone (white bars) or with the addition of 1 μg/mL EIX (black bars). Ethylene production was measured using gas-chromatography. Presented values are normalized to M82 mock average (NC). (a) Average ± SEM of 5 independent experiments is presented, N≥8. (b) Average ± SEM of 4 independent experiments is presented, N≥12. Asterisks represent statistical significance in a two tailed t-test (*p <0.05, **p<0.01, ***p<0.001, ****p<0.0001).

**Supplemental Figure 2.**
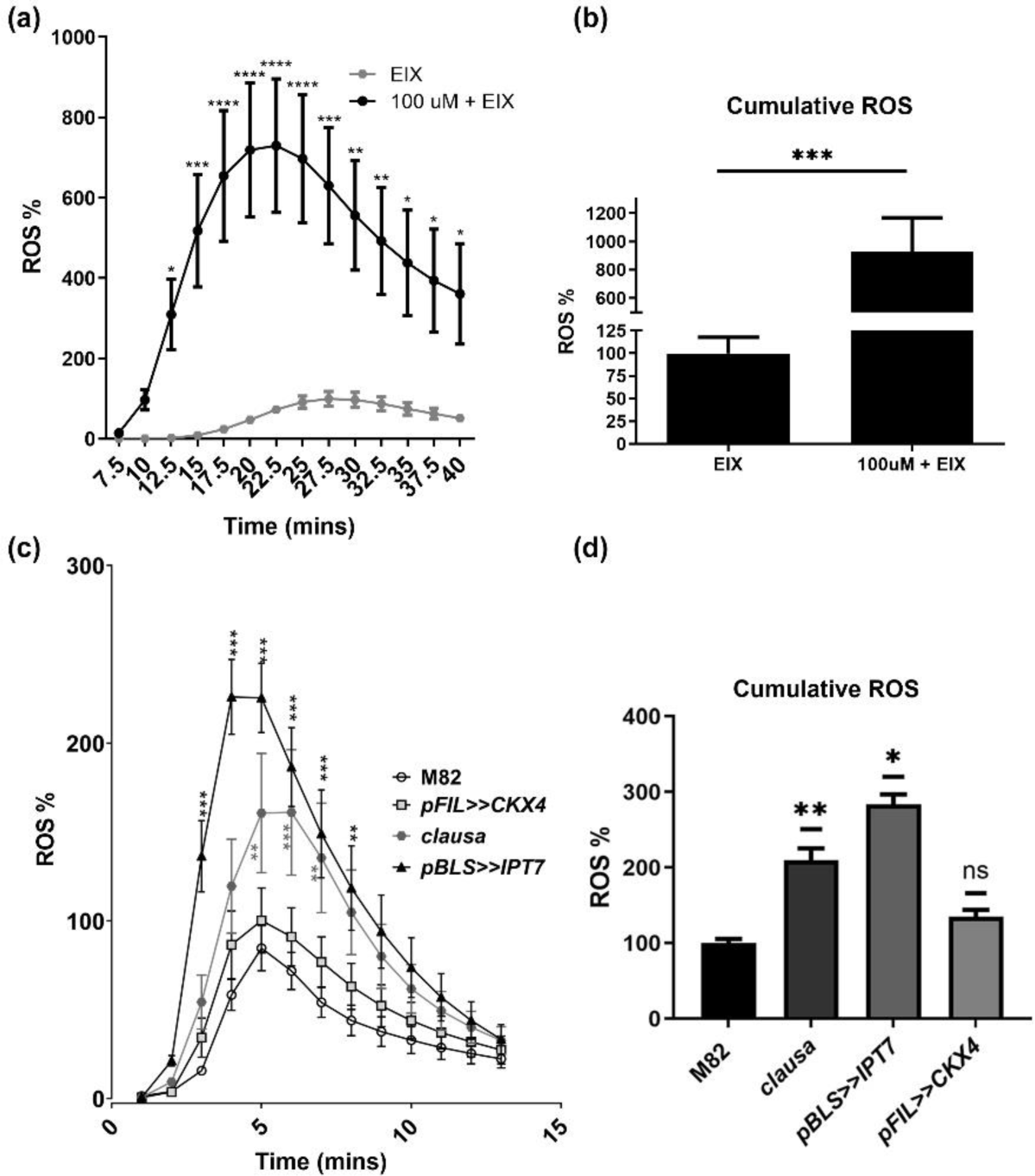
Cytokinin (CK) and Ethylene Inducing Xylanase (EIX) induce immune responses through separate pathways-ROS production. (a)-(b) *S. lycopersicum cv* M82 leaves were treated with 100uM 6-Benzylaminopurine (6-BAP) and 1 μg/mL EIX. (c)-(d) Leaves of the increased cytokinin (CK) transgene *pBLS>>IPT7*, the increased CK-sensitivity mutant *clausa e2522*, and the reduced CK content transgene *pFIL>>CKX4*, all in a *S. lycopersicum cv* M82 background, were treated with 1 μg/mL EIX. ROS production was measured every 5 minutes immediately after EIX application for 90 minutes using the HRP-luminol method. Average ± SEM of 4 independent experiments is shown, N=48. Asterisks represent statistical significance in a two-way ANOVA with a Bonferroni post hoc test, p<0.01. Inset shows total ROS production. *p<0.05; **p<0.01; ***p<0.001; ****p<0.0001.

**Supplemental Table 1.**
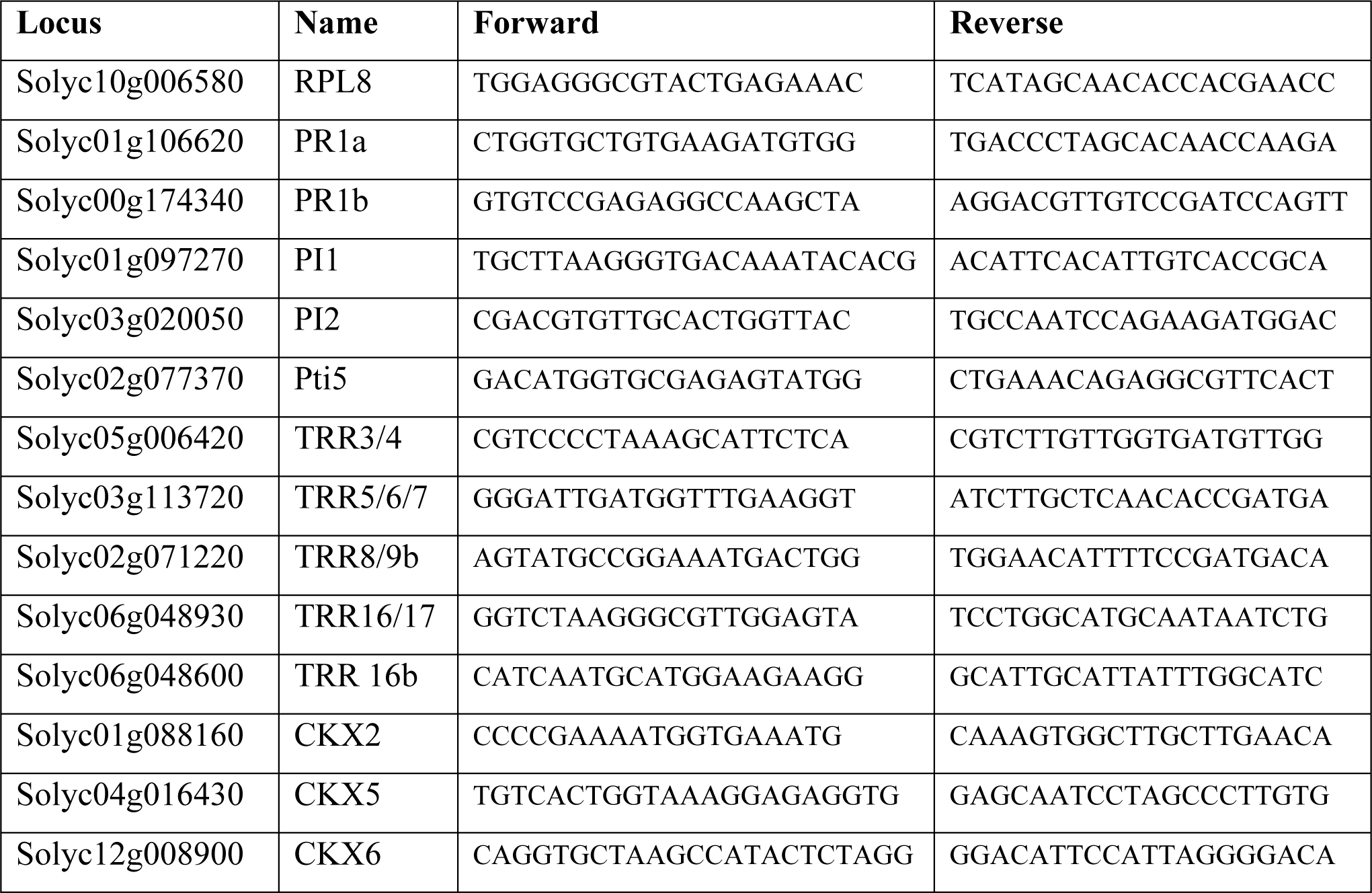
**qPCR primers.**

**Supplemental Table 2.**
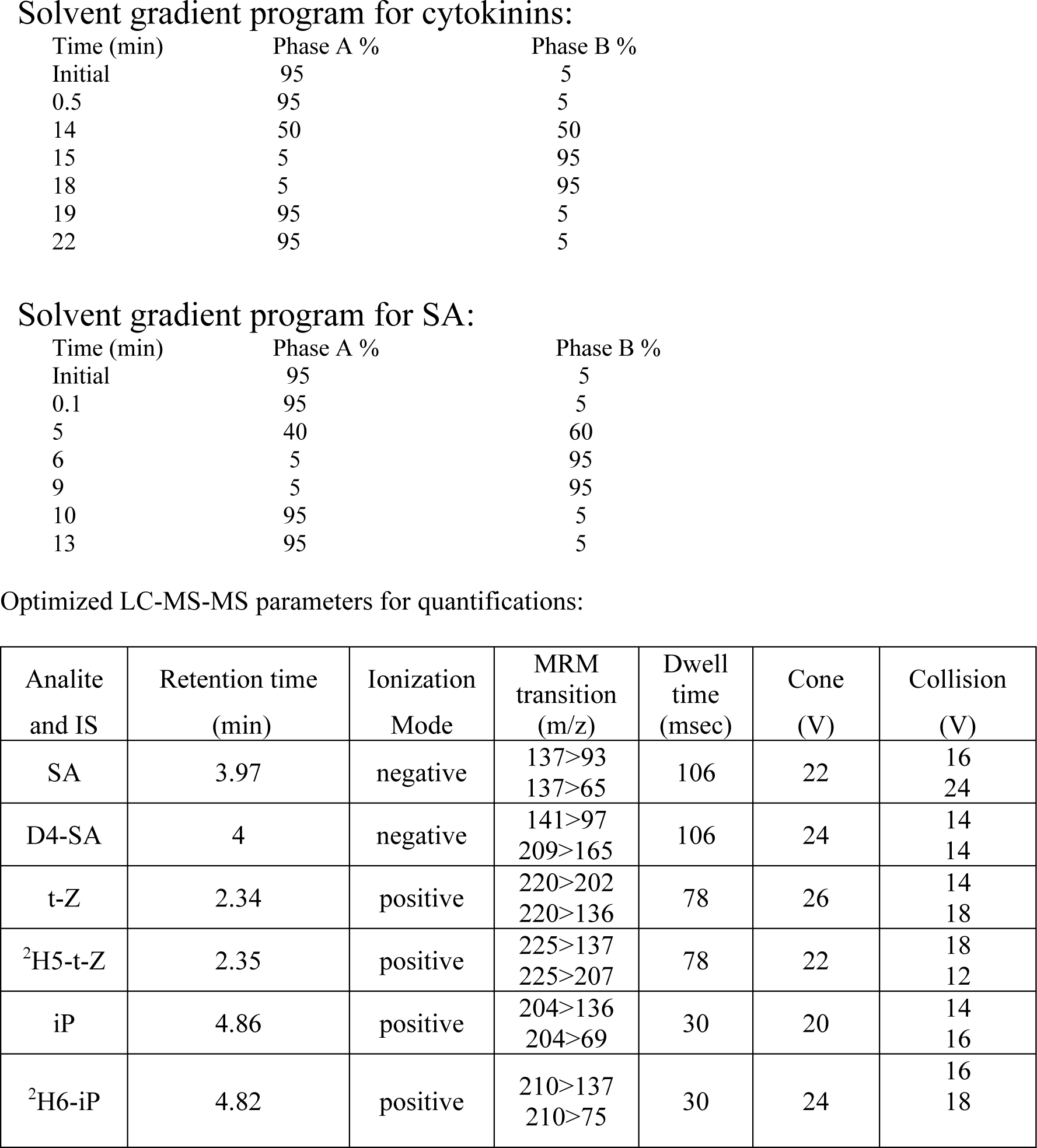
**Solvent gradients and MS-MS parameters for hormone quantification.**

## References

1. AbuQamar, S., Chai, M.F., Luo, H., Song, F.**, and** Mengiste, T. (2008). Tomato protein kinase 1b mediates signaling of plant responses to necrotrophic fungi and insect herbivory. Plant Cell 20: 1964–1983.

2. Albrecht, T. and Argueso, C.T. (2017). Should I fight or should I grow now? The role of cytokinins in plant growth and immunity and in the growth-defence trade-off. Annals of Botany 119: 725–735.

3. Ament, K., Kant, M.R., Sabelis, M.W., Haring, M.A.**, and** Schuurink, R.C. (2004). Jasmonic acid is a key regulator of spider mite-induced volatile terpenoid and methyl salicylate emission in tomato. Plant physiology 135: 2025–37.

4. Argueso, C.T., Ferreira, F.J., Epple, P., To, J.P.C., Hutchison, C.E., Schaller, G.E., Dangl, J.L.**, and** Kieber, J.J. (2012). Two-component elements mediate interactions between cytokinin and salicylic acid in plant immunity. PLoS Genetics 8.

5. Ballaré, C.L. (2011). Jasmonate-induced defenses: a tale of intelligence, collaborators and rascals. Trends in Plant Science 16: 249–257.

6. Bar, M. and Avni, A. (2009). EHD2 inhibits ligand-induced endocytosis and signaling of the leucine-rich repeat receptor-like protein LeEix2. The Plant Journal 59: 600–611.

7. Bar, M. and Avni, A. (2013). Endocytosis of LeEix and EHD proteins during plant defense signalling.

8. Bar, M., Israeli, A., Levy, M., Ben Gera, H., Jiménez-Gómez, J.M., Kouril, S., Tarkowski, P.**, and** Ori, N. (2016). CLAUSA Is a MYB Transcription Factor That Promotes Leaf Differentiation by Attenuating Cytokinin Signaling. The Plant cell 28: 1602–15.

9. Bari, R. and Jones, J.D.G. (2009). Role of plant hormones in plant defence responses. Plant molecular biology 69: 473–88.

10. Berens, M.L., Berry, H.M., Mine, A., Argueso, C.T.**, and** Tsuda, K. (2017). Evolution of Hormone Signaling Networks in Plant Defense. Annual Review of Phytopathology 55: 401–425.

11. Betsuyaku, S., Katou, S., Takebayashi, Y., Sakakibara, H., Nomura, N.**, and** Fukuda, H. (2018). Salicylic Acid and Jasmonic Acid Pathways are Activated in Spatially Different Domains Around the Infection Site During Effector-Triggered Immunity in Arabidopsis thaliana. Plant and Cell Physiology 59: 8–16.

12. Brading, P.A., Hammond-Kosack, K.E., Parr, A.**, and** Jones, J.D.G. (2000). Salicylic acid is not required for *Cf-2* - and *Cf-9* -dependent resistance of tomato to *Cladosporium fulvum*. The Plant Journal 23: 305–318.

13. Chanclud, E. and Morel, J.-B. (2016). Plant hormones: a fungal point of view. Molecular plant pathology 17: 1289–97.

14. Choi, J., Choi, D., Lee, S., Ryu, C.-M.**, and** Hwang, I. (2011). Cytokinins and plant immunity: old foes or new friends? Trends in Plant Science 16: 388–394.

15. Choi, J., Huh, S.U., Kojima, M., Sakakibara, H., Paek, K.-H.**, and** Hwang, I. (2010). The cytokinin-activated transcription factor ARR2 promotes plant immunity via TGA3/NPR1- dependent salicylic acid signaling in Arabidopsis. Developmental cell 19: 284–95.

16. Coenen, C. and Lomax, T.L. (1998). The diageotropica gene differentially affects auxin and cytokinin responses throughout development in tomato. Plant Physiology 117: 63–72.

17. Conrath, U., Beckers, G.J.M., Langenbach, C.J.G.**, and** Jaskiewicz, M.R. (2015). Priming for Enhanced Defense. Annual Review of Phytopathology 53: 97–119.

18. Cui, H., Sun, Y., Zhao, Z.**, and** Zhang, Y. (2019). The Combined Effect of Elevated O3 Levels and TYLCV Infection Increases the Fitness of Bemisia tabaci Mediterranean on Tomato Plants. Environmental Entomology.

19. D’haene, B., Vandesompele, J.**, and** Hellemans, J. (2010). Accurate and objective copy number profiling using real-time quantitative PCR. Methods 50: 262–270.

20. Dodds, P.N. and Rathjen, J.P. (2010). Plant immunity: towards an integrated view of plant– pathogen interactions. Nature Reviews Genetics 11: 539–548.

21. Durrant, W.E. and Dong, X. (2004). Systemic acquired resistance. Annual review of phytopathology 42: 185–209.

22. Elbaz, M., Avni, A.**, and** Weil, M. (2002). Constitutive caspase-like machinery executes programmed cell death in plant cells. Cell Death and Differentiation 9: 726–733.

23. Farber, M., Attia, Z.**, and** Weiss, D. (2016). Cytokinin activity increases stomatal density and transpiration rate in tomato. Journal of Experimental Botany 67: 6351–6362.

24. Gordon, S.P., Chickarmane, V.S., Ohno, C.**, and** Meyerowitz, E.M. (2009). Multiple feedback loops through cytokinin signaling control stem cell number within the Arabidopsis shoot meristem. Proceedings of the National Academy of Sciences 106: 16529–16534.

25. Grosskinsky, D.K. et al. (2011). Cytokinins Mediate Resistance against Pseudomonas syringae in Tobacco through Increased Antimicrobial Phytoalexin Synthesis Independent of Salicylic Acid Signaling. Plant Physiology 157: 815–830.

26. Harel, Y.M., Mehari, Z.H., Rav-David, D.**, and** Elad, Y. (2014). Systemic Resistance to Gray Mold Induced in Tomato by Benzothiadiazole and *Trichoderma harzianum* T39. Phytopathology 104: 150–157.

27. Hatsugai, N., Igarashi, D., Mase, K., Lu, Y., Tsuda, Y., Chakravarthy, S., Wei, H., Foley, J.W., Collmer, A., Glazebrook, J.**, and** Katagiri, F. (2017). A plant effector-triggered immunity signaling sector is inhibited by pattern-triggered immunity. The EMBO Journal 36: 2758–2769.

28. Iberkleid, I., Ozalvo, R., Feldman, L., Elbaz, M., Patricia, B.**, and** Horowitz, S.B. (2014). Responses of Tomato Genotypes to Avirulent and *Mi* -Virulent *Meloidogyne javanica* Isolates Occurring in Israel. Phytopathology 104: 484–496.

29. Jameson, P.E. (2000). Cytokinins and auxins in plant-pathogen interactions – An overview. Plant Growth Regulation 32: 369–380.

30. Jiang, C.-J.J., Shimono, M., Sugano, S., Kojima, M., Liu, X., Inoue, H., Sakakibara, H.**, and** Takatsuji, H. (2013). Cytokinins act synergistically with salicylic acid to activate defense gene expression in rice. Molecular Plant-Microbe Interactions 26: 287–296.

31. Jones, J.D.G. and Dangl, J.L. (2006). The plant immune system. Nature 444: 323–329.

32. Karasov, T.L., Chae, E., Herman, J.J.**, and** Bergelson, J. (2017). Mechanisms to Mitigate the Trade-Off between Growth and Defense. The Plant Cell 29: 666–680.

33. Keshishian, E.A. and Rashotte, A.M. (2015). Plant cytokinin signalling. Essays in biochemistry 58: 13–27.

34. Király, Z., El Hammady, M., Pozsár, B. and Kiraly, Z., Hammady, M.E.**, and** Pozsar, B. (1967). Increased cytokinin activity of rust-infected bean and broad bean leaves. Phytopathology 57: 93–94.

35. Kiraly, Z., Pozsar, B.**, and** Hammady, M.E. (1966). Cytokinin activity in rust infected plants: juvenility and senescence in diseased leaf tissues. Acta Phytopathol. Acad. Sci. Hung. 1: 29–37.

36. Kurakawa, T., Ueda, N., Maekawa, M., Kobayashi, K., Kojima, M., Nagato, Y., Sakakibara, H.**, and** Kyozuka, J. (2007). Direct control of shoot meristem activity by a cytokinin-activating enzyme. Nature 445: 652–655.

37. Lashbrook, C.C., Tieman, D.M.**, and** Klee, H.J. (1998). Differential regulation of the tomato ETR gene family throughout plant development. The Plant journal : for cell and molecular biology 15: 243–52.

38. Leibman-Markus, M., Schuster, S.**, and** Avni, A. (2017a). LeEIX2 Interactors’ Analysis and EIX-Mediated Responses Measurement. In Methods in molecular biology (Clifton, N.J.), pp. 167–172.

39. Leibman-Markus, M., Schuster, S.**, and** Avni, A. (2017b). LeEIX2 Interactors’ Analysis and EIX-Mediated Responses Measurement. In Plant Pattern Recognition Receptors: Methods and Protocols, L. Shan and P. He, eds (Springer New York: New York, NY), pp. 167–172.

40. Li, L., Li, C., Lee, G.I.**, and** Howe, G.A. (2002). Distinct roles for jasmonate synthesis and action in the systemic wound response of tomato. Proceedings of the National Academy of Sciences of the United States of America 99: 6416–21.

41. Li, Y., Qin, L., Zhao, J., Muhammad, T., Cao, H., Li, H., Zhang, Y., and Liang, Y. (2017). SlMAPK3 enhances tolerance to tomato yellow leaf curl virus (TYLCV) by regulating salicylic acid and jasmonic acid signaling in tomato (Solanum lycopersicum). PLOS ONE 12: e0172466.

42. Liu, L., Sonbol, F.-M., Huot, B., Gu, Y., Withers, J., Mwimba, M., Yao, J., He, S.Y.**, and** Dong, X. (2016). Salicylic acid receptors activate jasmonic acid signalling through a non-canonical pathway to promote effector-triggered immunity. Nature Communications 7.

43. López-Ráez, J.A., Verhage, A., Fernández, I., García, J.M., Azcón-Aguilar, C., Flors, V.**, and** Pozo, M.J. (2010). Hormonal and transcriptional profiles highlight common and differential host responses to arbuscular mycorrhizal fungi and the regulation of the oxylipin pathway. Journal of Experimental Botany 61: 2589–2601.

44. Macho, A.P. and Zipfel, C. (2014). Plant PRRs and the activation of innate immune signaling. Molecular cell 54: 263–72.

45. Marhavý, P., Bielach, A., Abas, L., Abuzeineh, A., Duclercq, J., Tanaka, H., Pařezová, M., Petrášek, J., Friml, J., Kleine-Vehn, J.**, and** Benková, E. (2011). Cytokinin Modulates Endocytic Trafficking of PIN1 Auxin Efflux Carrier to Control Plant Organogenesis. Developmental Cell 21: 796–804.

46. Martínez-Medina, A., Fernández, I., Sánchez-Guzmán, M.J., Jung, S.C., Pascual, J.A.**, and** Pozo, M.J. (2013). Deciphering the hormonal signalling network behind the systemic resistance induced by Trichoderma harzianum in tomato. Frontiers in Plant Science 4: 206.

47. Mehari, Z.H., Elad, Y., Rav-David, D., Graber, E.R.**, and** Meller Harel, Y. (2015). Induced systemic resistance in tomato (Solanum lycopersicum) against Botrytis cinerea by biochar amendment involves jasmonic acid signaling. Plant and Soil 395: 31–44.

48. Mok, D.W. and Mok, M.C. (2001). Cytokinin metabolism and action. Ann. Rev. Plant Physiol. Plant Mol. Biol. 52: 89–118.

49. Muller, B. and Sheen, J. (2007). Arabidopsis Cytokinin Signaling Pathway. Science’s STKE 2007: cm5-cm5.

50. Naseem, M. and Dandekar, T. (2012). The Role of Auxin-Cytokinin Antagonism in Plant-Pathogen Interactions. PLoS Pathogens 8.

51. Naseem, M., Philippi, N., Hussain, A., Wangorsch, G., Ahmed, N., Dandekara, T.**, and** Dandekar, T. (2012). Integrated systems view on Networking by hormones in Arabidopsis immunity reveals multiple crosstalk for cytokinin. Plant Cell 24: 1793–1814.

52. Pertry, I. et al. (2009). Identification of Rhodococcus fascians cytokinins and their modus operandi to reshape the plant. Proceedings of the National Academy of Sciences of the United States of America 106: 929–34.

53. Pizarro, L., Leibman-Markus, M., Schuster, S., Bar, M., Meltz, T.**, and** Avni, A. (2018). Tomato Prenylated RAB Acceptor Protein 1 Modulates Trafficking and Degradation of the Pattern Recognition Receptor LeEIX2, Affecting the Innate Immune Response. Frontiers in Plant Science 9: 257.

54. Pogány, M., Koehl, J., Heiser, I., Elstner, E.F.**, and** Barna, B. (2004). Juvenility of tobacco induced by cytokinin gene introduction decreases susceptibility to Tobacco necrosis virus and confers tolerance to oxidative stress. Physiological and Molecular Plant Pathology 65: 39–47.

55. Robert-Seilaniantz, A., Grant, M.**, and** Jones, J.D.G.G. (2011). Hormone crosstalk in plant disease and defense: more than just jasmonate-salicylate antagonism. Annual review of phytopathology 49: 317–43.

56. Romanov, G.A., Lomin, S.N.**, and** Schmülling, T. (2006). Biochemical characteristics and ligand-binding properties of Arabidopsis cytokinin receptor AHK3 compared to CRE1/AHK4 as revealed by a direct binding assay. Journal of Experimental Botany 57: 4051–4058.

57. Ron, M. and Avni, A. The Receptor for the Fungal Elicitor Ethylene-Inducing Xylanase Is a Member of a Resistance-Like Gene Family in Tomato.

58. Ron, M., Kantety, R., Martin, G.B., Avidan, N., Eshed, Y., Zamir, D., and Avni, A. (2000). High-resolution linkage analysis and physical characterization of the EIX-responding locus in tomato. Theor. Appl. Genet. 100: 184–189.

59. Ryals, J.A., Neuenschwander, U.H., Willits, M.G., Molina, A., Steiner, H.Y.**, and** Hunt, M.D. (1996). Systemic Acquired Resistance. The Plant cell 8: 1809–1819.

60. Sakakibara, H. (2006). Cytokinins: Activity, Biosynthesis, and Translocation. Annu Rev Plant Biol.

61. Schindelin, J. et al. (2012). Fiji: an open-source platform for biological-image analysis. Nature Methods 9: 676–682.

62. Shani, E., Ben-Gera, H., Shleizer-Burko, S., Burko, Y., Weiss, D.**, and** Ori, N. (2010). Cytokinin Regulates Compound Leaf Development in Tomato. Plant Cell 22: 3206– 3217.

63. Sharfman, M., Bar, M., Ehrlich, M., Schuster, S., Melech-Bonfil, S., Ezer, R., Sessa, G.**, and** Avni, A. (2011). Endosomal signaling of the tomato leucine-rich repeat receptor-like protein LeEix2. The Plant Journal 68: 413–423.

64. Sharon, A., Fuchs, Y.**, and** Anderson, J.D. (1993). The Elicitation of Ethylene Biosynthesis by a Trichoderma Xylanase Is Not Related to the Cell Wall Degradation Activity of the Enzyme. Plant physiology 102: 1325–1329.

65. Shaya, F., David, I., Yitzhak, Y.**, and** Izhaki, A. (2019). Hormonal interactions during early physiological partenocarpic fruitlet abscission in persimmon (Diospyros Kaki Thunb.) ‘Triumph’ and ‘Shinshu’ cultivars. Scientia Horticulturae 243: 575–582.

66. Shigenaga, A.M., Berens, M.L., Tsuda, K.**, and** Argueso, C.T. (2017). Towards engineering of hormonal crosstalk in plant immunity. Current Opinion in Plant Biology 38: 164–172.

67. Shoresh, M., Yedidia, I.**, and** Chet, I. (2005). Involvement of Jasmonic Acid/Ethylene Signaling Pathway in the Systemic Resistance Induced in Cucumber by *Trichoderma asperellum* T203. Phytopathology 95: 76–84.

68. Spoel, S.H. and Dong, X. (2012). How do plants achieve immunity? Defence without specialized immune cells. Nature reviews. Immunology 12: 89–100.

69. Thara, V.K., Tang, X., Gu, Y.Q., Martin, G.B.**, and** Zhou, J.-M. (1999). Pseudomonas syringae pv tomato induces the expression of tomato EREBP-like genes Pti4 and Pti5 independent of ethylene, salicylate and jasmonate. The Plant Journal 20: 475–483.

70. Thomma, B.P., Eggermont, K., Penninckx, I.A., Mauch-Mani, B., Vogelsang, R., Cammue, B.P.**, and** Broekaert, W.F. (1998). Separate jasmonate-dependent and salicylate-dependent defense-response pathways in Arabidopsis are essential for resistance to distinct microbial pathogens. Proceedings of the National Academy of Sciences of the United States of America 95: 15107–11.

71. Werner, T. and Schmülling, T. (2009). Cytokinin action in plant development. Current opinion in plant biology 12: 527–38.

72. Zdarska, M., Dobisová, T., Gelová, Z., Pernisová, M., Dabravolski, S.**, and** Hejátko, J. (2015). Illuminating light, cytokinin, and ethylene signalling crosstalk in plant development. Journal of experimental botany 66: 4913–31.

73. Zürcher, E., Tavor-Deslex, D., Lituiev, D., Enkerli, K., Tarr, P.T.**, and** Müller, B. (2013). A robust and sensitive synthetic sensor to monitor the transcriptional output of the cytokinin signaling network in planta. Plant physiology 161: 1066–75.

